# Affinity-engineered human antibodies detect celiac disease gluten pMHC complexes and inhibit T-cell activation

**DOI:** 10.1101/840561

**Authors:** Rahel Frick, Lene S. Høydahl, Ina Hodnebrug, Shraddha Kumari, Grete Berntsen, Jeliazko R. Jeliazkov, Kristin S. Gunnarsen, Terje Frigstad, Erik S. Vik, Knut E.A. Lundin, Sheraz Yaqub, Jørgen Jahnsen, Jeffrey J. Gray, Ludvig M. Sollid, Inger Sandlie, Geir Åge Løset

## Abstract

Antibodies specific for antigenic peptides bound to major histocompatibility complex (MHC) molecules are valuable tools for studies of antigen presentation. Such T-cell receptor (TCR)-like antibodies may also have therapeutic potential in human disease due to their ability to target disease-associated antigens with high specificity. We previously generated celiac disease (CeD) relevant TCR-like antibodies that recognize the prevalent gluten epitope DQ2.5-glia-α1a in complex with HLA-DQ2.5. Here, we report on second-generation high-affinity antibodies towards this epitope as well as a panel of novel TCR-like antibodies to another immunodominant gliadin epitope, DQ2.5-glia-α2. The strategy for affinity engineering was based on Rosetta modeling combined with pIX phage display and is applicable to similar protein engineering efforts. We isolated picomolar affinity binders and validated them in Fab and IgG format. Flow cytometry experiments with CeD biopsy material confirm the unique disease specificity of these TCR-like antibodies and reinforce the notion that B cells and plasma cells have a dominant role in gluten antigen presentation in the inflamed CeD gut. Further, the lead candidate 3.C11 potently inhibited CD4^+^ T-cell activation and proliferation *in vitro* in an HLA and epitope specific manner, pointing to a potential for targeted disease interception without compromising systemic immunity.

**Significance Statement:** Consumption of gluten-containing food drives celiac disease in genetically predisposed individuals. The underlying disease mechanism is not fully understood, but it is strictly dependent on activation of pathogenic T cells. We have engineered high-affinity human antibodies recognizing the T-cell target HLA-DQ2.5 in complex with gluten epitopes and studied cell-specific antigen presentation in patients, which shows that plasma cells and not dendritic cells dominate the inflamed tissue. The only available treatment is lifelong adherence to a gluten-free diet, which is difficult and not effective in all cases. We show that at least one of our antibodies can specifically inhibit activation of pathogenic T-cells *in vitro* and therefore shows promise for therapy.

## Introduction

T-cell receptors (TCRs) and B-cell receptors (BCRs) are the two classes of antigen-specific receptors on cells of the adaptive immune system. While TCRs recognize peptide fragments of antigen in the context of cell-surface major histocompatibility complex (MHC) class I or class II proteins, BCRs and their secreted counterparts, antibodies, bind different soluble or cell-surface antigens without requirements for scaffold proteins (1). Soluble reagents with specificity for peptide:MHC (pMHC) complexes have been engineered as tools to study antigen presentation and as potential therapeutics in cancer and autoimmune disease (2, 3). While membrane bound TCRs are the natural binding partners for pMHC, soluble TCRs require extensive stability and affinity engineering before they can be used as detection reagents (4, 5). Antibodies are more stable, have higher affinities and can be readily expressed as soluble molecules (6). Their modular architecture, consisting of antigen binding (Fab) and crystallizable (Fc) fragments, facilitates their use as detection reagents and adds effector functions. Generating human antibodies with TCR-like specificity is challenging. Nevertheless, TCR-like antibodies specific for MHC class I or class II complexes have been generated by immunization strategies and the use of *in vitro* display technologies, most commonly phage display (2). These antibodies have been successfully used to detect and quantify peptide presentation on cells and several studies suggest therapeutic potential by different modes of action, including inhibition of pathogenic T cells and killing mechanisms to delete antigen presenting cells (APCs) (7–9).

In this study, we engineered high affinity antibodies specific for pMHC complexes implicated in celiac disease (CeD). CeD is an inflammatory autoimmune-like condition of the small intestine caused by immune reactions to dietary gluten proteins. The disease is driven by CD4^+^ T cells that recognize deamidated gluten peptides in the context of disease-associated HLA-DQ molecules, chiefly HLA-DQ2.5 (*DQA1*05 – DQB1*02*) (10). The deamidation of gluten peptides is mediated by the enzyme transglutaminase 2 (TG2) and results in conversion of Gln to negatively charged Glu at specific sites in polypeptides. This posttranslational modification is a key event transforming proteolytically stable, but immunologically inert, peptides into pathogenic T-cell epitopes (11, 12). A range of gluten T-cell epitopes has been characterized, but four immunodominant epitopes derived from α-gliadin and ω-gliadin are particularly prominent, namely DQ2.5-glia-α1a (PFPQPELPY), DQ2.5-glia-α2 (PQPELPYPQ), DQ2.5-glia-ω1 (PFPQPEQPF), and DQ2.5-glia-ω2 (PQPEQPFPW). T-cell responses against these epitopes are found in the majority of patients and are believed to orchestrate tissue destruction in the small intestine and autoantibody production (13–15). The only currently available treatment for CeD is lifelong adherence to a gluten-free diet. Development of alternative treatments is sought after due to poor patient compliance with the dietary restrictions, unavailability of strictly gluten free food and the feared transformation of uncomplicated CeD to refractory CeD (16, 17).

Recently, we reported the generation and use of antibodies selected on HLA-DQ2.5:DQ2.5-glia-α1a. We surprisingly found that gut plasma cells (PCs) are the most abundant gliadin peptide-presenting cells in the inflamed small intestine of CeD patients and that they express both HLA class II and T cell co-stimulatory molecules (18).

In the current study, we describe the generation and affinity maturation of human TCR-like antibodies specific for HLA-DQ2.5 in complex with either DQ2.5-glia-α1a or DQ2.5-glia-α2. Our strategy was based on structural modeling using the Rosetta software suite in combination with pIX phage display and gave rise to highly specific binders with picomolar monomeric affinities. The antibodies stained PCs from inflamed CeD patient biopsy material, whereas the scarce CD11c^+^ and CD14^+^ dendritic cells (DCs) and macrophages (Mfs) stained negligibly. This confirms and extends our previous observation on the central role of the B-cell compartment in this tissue (18). We then chose the high-affinity TCR-like antibody with specificity towards the immunodominant DQ2.5-glia-α2 epitope for *in vitro* studies of inhibition of T-cell activation. Potent and strictly HLA and epitope dependent T-cell inhibition was observed *in vitro*, suggesting that this antibody has potential to be used for disease-specific immunotherapy.

## Results

### Selection of an antibody specific for HLA-DQ2.5 with bound DQ2.5-glia-α2

To characterize the APC subsets involved in gluten peptide presentation in CeD patients, we previously generated an antibody specific for HLA-DQ2.5:DQ2.5-glia-α1a by phage display (18). Here we extend the study to include an antibody with specificity for another dominant epitope derived from α-gliadin, DQ2.5-glia-α2, in complex with HLA-DQ2.5. To this end, a human naïve scFv phage display library (19) was panned against soluble, recombinant pMHC. After three rounds of selection, we assessed antigen reactivity of the selection output in a polyclonal ELISA and observed increased and preferential binding to the target (S. 1A). We then reformatted the selection output from the phagemid to a vector for soluble scFv expression and screened 70 single clones for target binding by ELISA (**Fig. 1**A). A total of 49 single clones bound target preferentially, and sequence analysis identified 14 unique sequences (**Fig. 1**A and S. 1B). Five of the clones were enriched in the selection output. To characterize the selected scFvs and choose a lead clone, we expressed all unique clones in *E. coli* and directly compared periplasmic fractions in ELISA for target binding (**Fig. 1**B). Next, we performed crude SPR measurements using purified scFv (S. 1C). Based on reactivity profiles and apparent affinities, we chose a lead clone, termed 206. When reformatted to full-length human IgG1 (hIgG1), 206 retained binding to HLA-DQ2.5:DQ2.5-glia-α2, albeit weakly (**Fig. 1**C, S. 1C). No binding to the highly similar epitope HLA-DQ2.5:DQ2.5-glia-ω2 was observed (**Fig. 1**D). To accurately determine the monomeric affinity, a Fab version of 206 was constructed and SPR analysis estimated the monomeric affinity to K_D_ 240±20 nM with a high off-rate (2.4×10^−1^ s^−1^) (**Fig. 1**E). The previously characterized lead antibody specific for HLA-DQ2.5:DQ2.5-glia-α1a, termed 107, was determined to have a dissociation constant (K_D_) of approximately 70 nM (18).

**Fig. 1:**
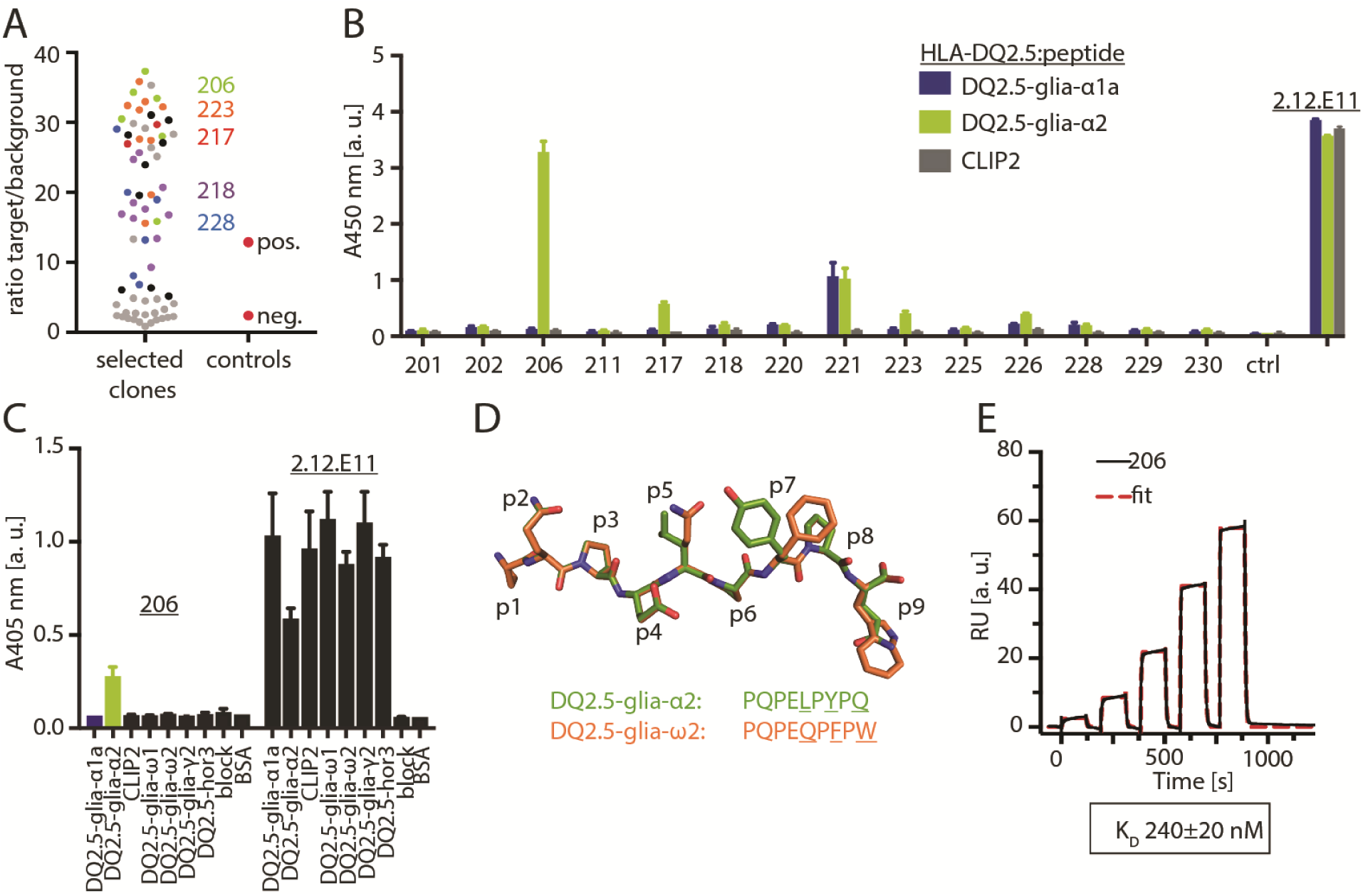
Isolation and characterization of an antibody specific for HLA-DQ2.5:DQ2.5-glia-α2. **A:** The selection output after three rounds of selection was batch cloned into a vector for soluble scFv expression and single colonies were expressed in the *E. coli* periplasm followed by screening for binding to HLA-DQ2.5:DQ2.5-glia-α2 (target) and HLA-DQ2.5:CLIP2 (background) in ELISA. ScFv anti-phOx binding to BSA-phOx and empty *E. coli* were included as positive and negative controls, respectively. The ratios of target/background binding are represented as dots for each clone. Positive clones were sequenced, and colors represent clones found repeatedly. Grey dots denote clones with unknown sequence. **B:** 14 unique clones were expressed, and periplasmic fractions were analyzed for binding to related gliadin pMHC complexes in ELISA. *E. coli* XL1-Blue were included as a negative control (ctrl) and pMHC capture levels were controlled with the HLA-DQ2 antibody 2.12.E11. Error bars illustrate mean ± SD of duplicates (n=2). **C:** The lead candidate 206 was reformatted to hIgG1 and binding to a larger peptide panel of pMHC complexes was tested in ELISA. Error bars illustrate mean ± SD of duplicates (n=2). **D:** Structural alignment of DQ2.5-glia-α2 (PDB code 4OZH (20)) and the related DQ2.5-glia-ω2 epitope (model). Differing positions are underlined. **E:** The monomeric affinity of Fab 206 for HLA-DQ2.5:DQ2.5-glia-α2 was determined by SPR using single cycle kinetics and fitting a 1:1 binding model to the data (n=2).

### Structural models of the pMHC-specific antibodies and library design

In order to increase the affinity of both antibodies (107 and 206), we generated structural models of the antibody Fv domains using RosettaAntibody and predicted their interaction with the corresponding HLA-DQ2.5 complexes using SnugDock (18). Final models were selected manually from a set of 1,000 docked complexes generated for each antibody. Criteria were a low Rosetta interface score, concurrence with experimental specificity data and similarity to other low-scoring models (S. 2A+B). Overall, both 107 and 206 were predicted to bind pMHC in a diagonal TCR-like manner, with the footprint focused on the C-terminal part of the peptides (**Fig. 2**A, S. 2C+D). The light chains of both antibodies were predicted to have a number of favorable interactions with the MHC. Interestingly, relatively large parts of the CDR L1 and L3 loops have similar sequences in the two antibodies, and D28 and S30 of L1 and N92, S93, and Y94 of L3 were predicted to interact with the MHC α1 and β1 helices in a similar fashion (**Fig. 2**B). For both antibodies the models suggested that contacts between the CDR H1 and H3 loops and the pMHC could be introduced to increase the binding interface (**Fig. 2**C+D). The CDR H2 loops in contrast, were oriented away from the peptide and therefore seemed less promising targets for affinity maturation. In particular, the CDR H1 loops appeared to be too short to contact the peptide or MHC. We therefore generated focused libraries with sequence randomization and increased lengths of either the CDR H1 or H3 loop (Supplementary Table 1). As the low-scoring models of 107 positioned a tryptophan (W100) in a pocket between the peptide and the MHC binding groove, we retained this residue in 50% of the 107 H3 library clones. We achieved 1×10^9^ – 5×10^9^ primary transformants for each of the four CDR-targeted sub-libraries. Additionally, as paratope-distal residues may improve biophysical properties and affect affinity (21–23), we generated a random library with mutations across the entire scFv sequence of 107 by PCR using dNTP analogs, achieving an average amino acid mutagenesis load of 4% and 5×10^7^ primary transformants.

**Fig. 2:**
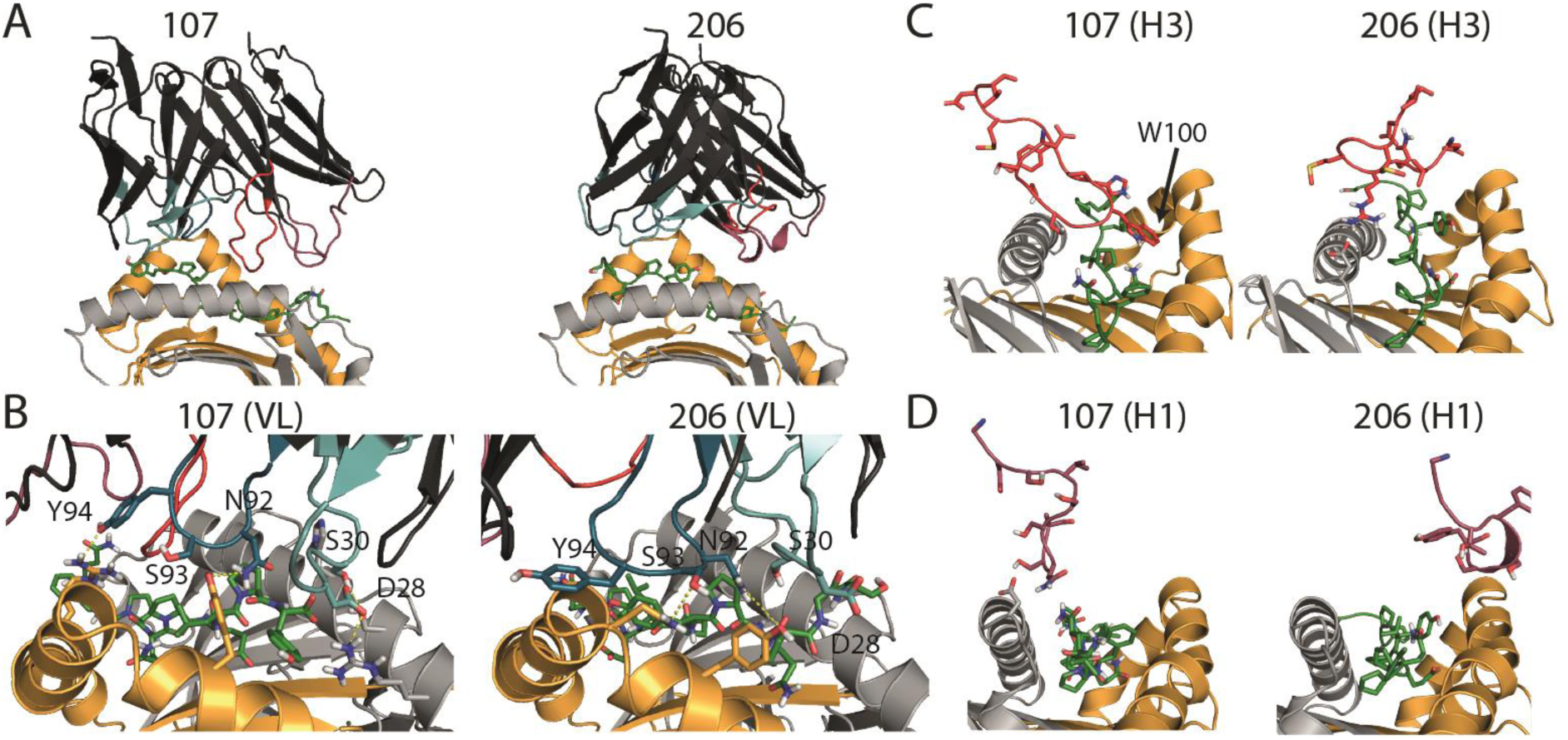
Fv-pMHC models and library design. **A:** Low-scoring models suggest a diagonal binding mode across the pMHC antigen-binding groove for antibodies 107 and 206. **B:** Residues in the light chain CDR loops that are conserved across the two antibodies and predicted to be within 3 Å of the pMHC are represented as sticks and labeled. **C+D:** The CDR H3 (C) and H1 (D) loops of 107 and 206 in cartoon representation with side chains as sticks. The buried W100 in H3 of 107 is labeled. Yellow: MHC β-chain, light grey: MHC α-chain, green: gliadin peptide, black: Fv frameworks, red: CDR loops of the heavy chain, teal: CDR loops of the light chain.

### Selection of high-affinity gluten pMHC-specific antibodies

For the selection of second-generation antibodies, we designed a phage display selection strategy aimed at isolating high-affinity antibodies with retained specificity and improved thermostability (**Fig. 3**A). All libraries were packaged at high valence (HV) display on phage coat protein pIX (24). After an initial low-stringency round (R1), the libraries were split into a thermostability branch and a competition branch for a highly stringent R2 with low target concentration followed by a non-stringent R3 loosely based on the hammer-hug selection protocol (25). In R2 of the competition branch, scFvs were displayed at low valence (LV). In the thermal branch, scFvs were displayed at HV and subjected to a heat challenge at temperatures that induced unfolding of the parent clones (S. 3). This was done prior to selection to aggregate and remove unstable library members (4, 26). In R3, we aimed at recovering and amplifying selected binders and therefore increased antigen concentration. For the thermal branch, we included a second heat challenge and displayed scFvs at LV to favor clones with high monomeric affinities. The stringent competition in R2 of the competition branch resulted in close to no selection output in all libraries, except for the 206 CDR H1 library. Libraries, from which no binders were retrieved, were discontinued. The thermostability branch however, yielded selection output from all libraries. To determine antigen reactivity of the outputs, we performed a polyclonal phage ELISA (S. 4A+B). Indeed, several libraries showed signs of enrichment of binders, and we continued to screen single clones from the R3 output as soluble scFv (**Fig. 3**B) and as scFv displayed on phage (**Fig. 3**C) by ELISA. Clones with preferential binding to the target were present in all libraries. Generally, the DQ2.5-glia-α2 binders bound with larger target/background ratios than the DQ2.5-glia-α1a binders. Several clones from the 107 random library bound modestly as scFv and displayed on phage, while the 107 CDR library clones only gave high signals when displayed on phage. Sequencing 50 offspring of antibody 107, revealed 35 unique DNA sequences and 17 unique amino acid sequences (S. 4C). All of these clones came from the CDR H3 library and possessed the central W100 residue, confirming its importance for binding. Sequencing the 206-derived selection output revealed that 66 out of 73 were unique DNA sequences, and 64 were unique at the amino acid level. Most of the unique DNA sequences (56/66) were derived from the CDR H1 library with a length increased by 2 residues, with the remaining clones (10/66) stemming from the CDR H1 pool with a length increase of 3 residues. Thus, there was only little clonal enrichment when comparing DNA sequences of selected binders, but distinct phenotypes appeared to be selected.

**Fig. 3:**
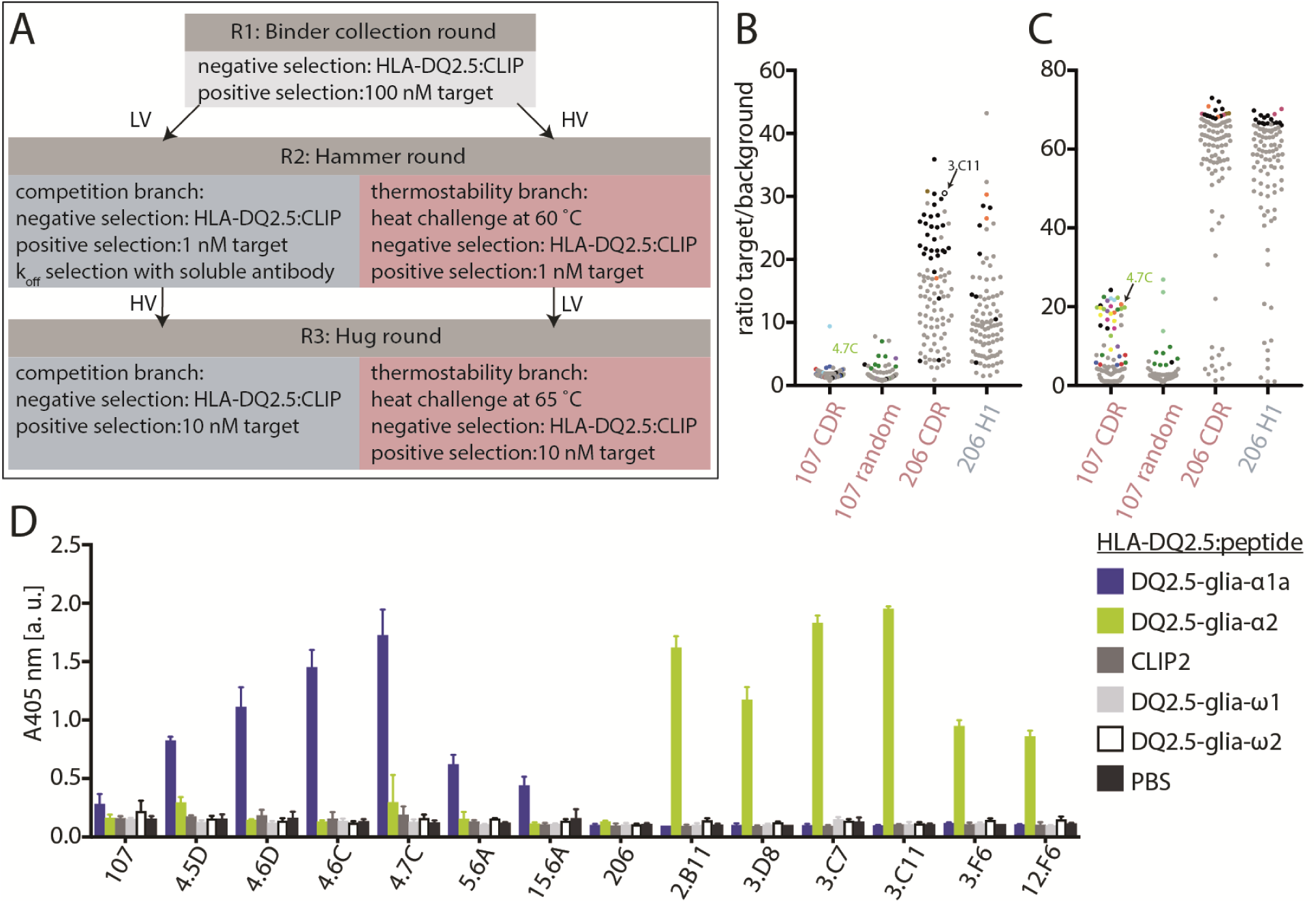
Selection and screening of antibody libraries. **A:** Schematic overview of the selection strategy. Low valence (LV) and high valence (HV) display were achieved by packaging with the helper phages M13KO7 or DeltaPhage, respectively. After R1 the libraries were split into a competition branch and a thermostability branch. **B:** The selection outputs after R3 were screened as soluble scFvs to assess binding to target pMHC complexes and HLA-DQ2.5:CLIP2 (background) in ELISA and ratios were calculated. Each dot represents one clone. Grey dots denote unknown sequences, black dots denote unique single sequences and colors represent enriched amino acid sequences. The 107 CDR, 107 random, and 206 CDR libraries were selected in the thermostability branch, while the 206 H1 library was selected in the competition branch. **C:** The selection outputs after R3 were screened in phage format and are represented as in B. **D:** Purified and monomeric Fab fragments were tested for binding to a larger panel of HLA-DQ2.5:peptide complexes in ELISA. pMHC levels were controlled with the HLA-DQ2 antibody 2.12.E11 (S. 4D). Error bars illustrate mean ± SD of duplicates (n=2).

Based on target binding in screening ELISAs and enrichment of sequence features, we chose 6 clones from each of the 107 and 206 outputs for large-scale Fab expression in HEK293E cells. These 12 clones were analyzed regarding their peptide-specificity in ELISA (**Fig. 3**D, S. 4D), and all were found to bind their target specifically. The DQ2.5-glia-α1a-specific variants did not cross react to the highly similar DQ2.5-glia-ω1 complex, which differs at only two positions in the peptide (p7 and p9) (S. 4E). Similarly, the DQ2.5-glia-α2-specific variants did not cross-react to the DQ2.5-glia-ω2 complex, which differs at three positions (p5, p7, and p9).

### Biophysical characterization of affinity matured pMHC-specific antibodies

We then performed binding analysis by SPR and ranked the 12 antibodies based on their off-rates (**Fig. 4**A and Supplementary Table 2). Strongly reduced off-rates were observed for all clones tested. The two 107-derived clones 5.6A and 15.6A, both from the random mutagenesis library, had less pronounced improvements than the four CDR H3 mutants selected from the library using structural models, underscoring the additional benefit of the targeted approach. Based on the results, we chose two clones, namely 4.7C and 3.C11, as leads for binding to HLA-DQ2.5:DQ2.5-glia-α1a and HLA-DQ2.5:DQ2.5-glia-α2, respectively.

**Fig. 4:**
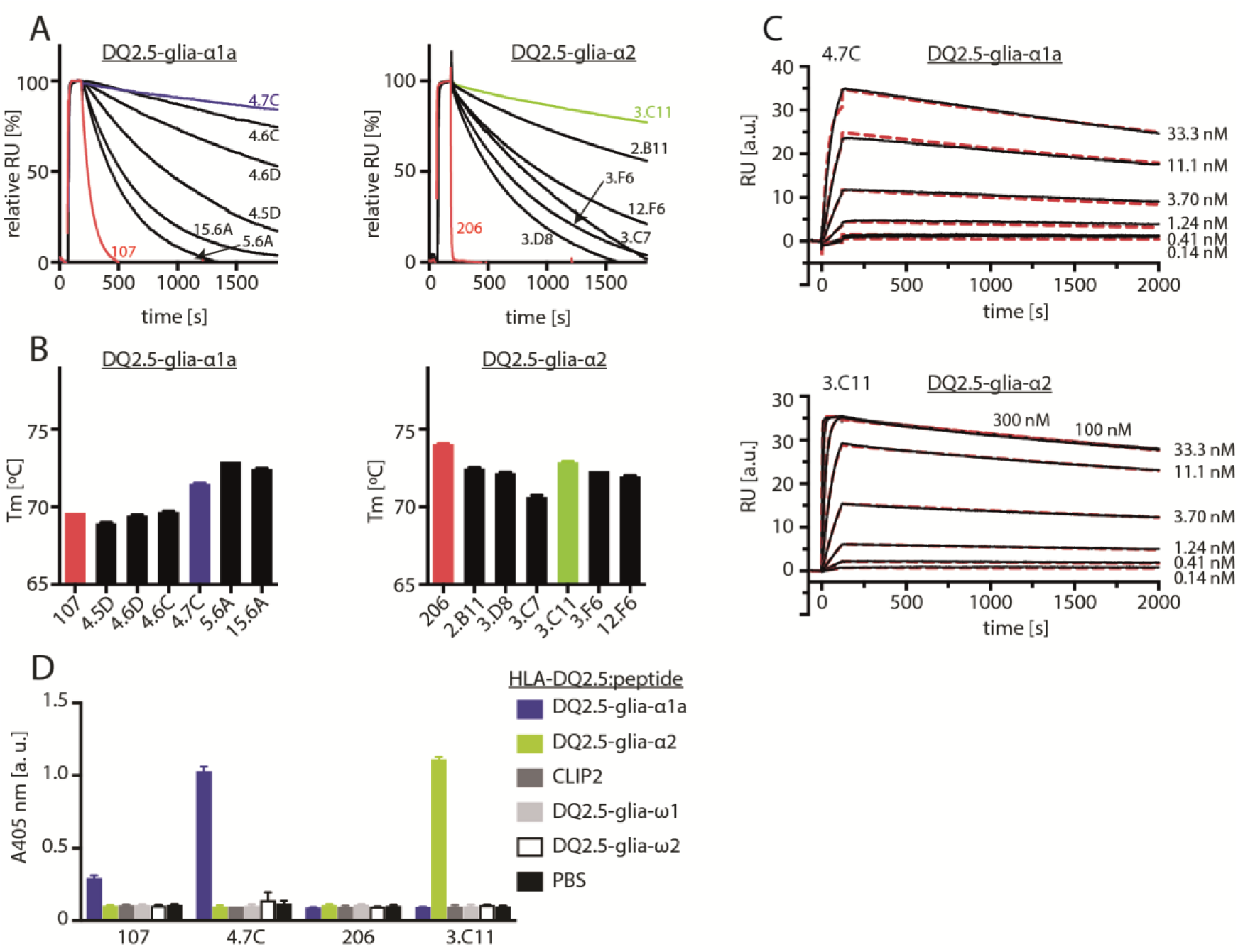
Biophysical characterization of leads. **A:** Fab fragments were ranked based on off-rates in SPR with the lead clones highlighted in blue and green and the first-generation antibodies in red. Left: binding to HLA-DQ2.5:DQ2.5-glia-α1a, Right: binding to HLA-DQ2.5:DQ2.5-glia-α2. **B:** Melting temperatures of the mother clones (red) and the affinity matured Fab fragments with the lead candidates highlighted in blue and green. Error bars illustrate mean ± SD of triplicates. **C:** Representative sensorgrams of 4.7C and 3.C11 (n≥ 2). **D:** The leads were reformatted to full-length hIgG1 and analyzed in ELISA against a panel of related soluble peptide:HLA-DQ2.5 complexes. Error bars illustrate mean ± SD of duplicates.

We next assessed the thermostability of all Fab fragments by determining their melting temperatures by nanoDSF (**Fig. 4**B). While most HLA-DQ2.5:DQ2.5-glia-α1a binders had improved thermostabilities compared to 107, the HLA-DQ2.5:DQ2.5-glia-α2 binders surprisingly had slightly lower melting temperatures than their mother clone. The lead clones, 4.7C and 3.C11, had the highest thermostabilities out of the binders isolated from targeted libraries. In line with the rational for generating the random mutagenesis libraries, the mutants 5.6A and 15.6A had the highest improvements in thermostability.

In concordance with the improved (lower) off-rates, all candidates had a strong improvement in affinity, with 4.7C, and 3.C11 having K_D_s of 170±40 pM and 88±8 pM, respectively (**Fig. 4**C and Supplementary Table 2). This is a 400-fold improvement for 4.7C and a 2,700-fold improvement for 3.C11. The second-generation antibodies were then expressed as full-length hIgG1 and tested for specific binding in ELISA (**Fig. 4**D). In agreement with previous results, 4.7C bound exclusively to the target complex HLA-DQ2.5:DQ2.5-glia-α1a, and 3.C11 to HLA-DQ2.5:DQ2.5-glia-α2. Thus, the high affinity antibodies maintained the high specificity of the parent clones.

### Structural basis for improved affinity

To understand the molecular interactions giving rise to improved affinity, Rosetta models of the two leads, 4.7C and 3.C11, were generated (**Fig. 5**A+B, S. 5). For 4.7C, several low-scoring models were highly similar to each other and to the model for the mother clone and were therefore selected as the final model. For 3.C11, two highly dissimilar docking modes were suggested by low-scoring models. Conservatively, the orientation most similar to the mother clone model was selected, as a strongly altered docking mode, with altered light-chain–pMHC contacts, is unlikely.

**Fig. 5:**
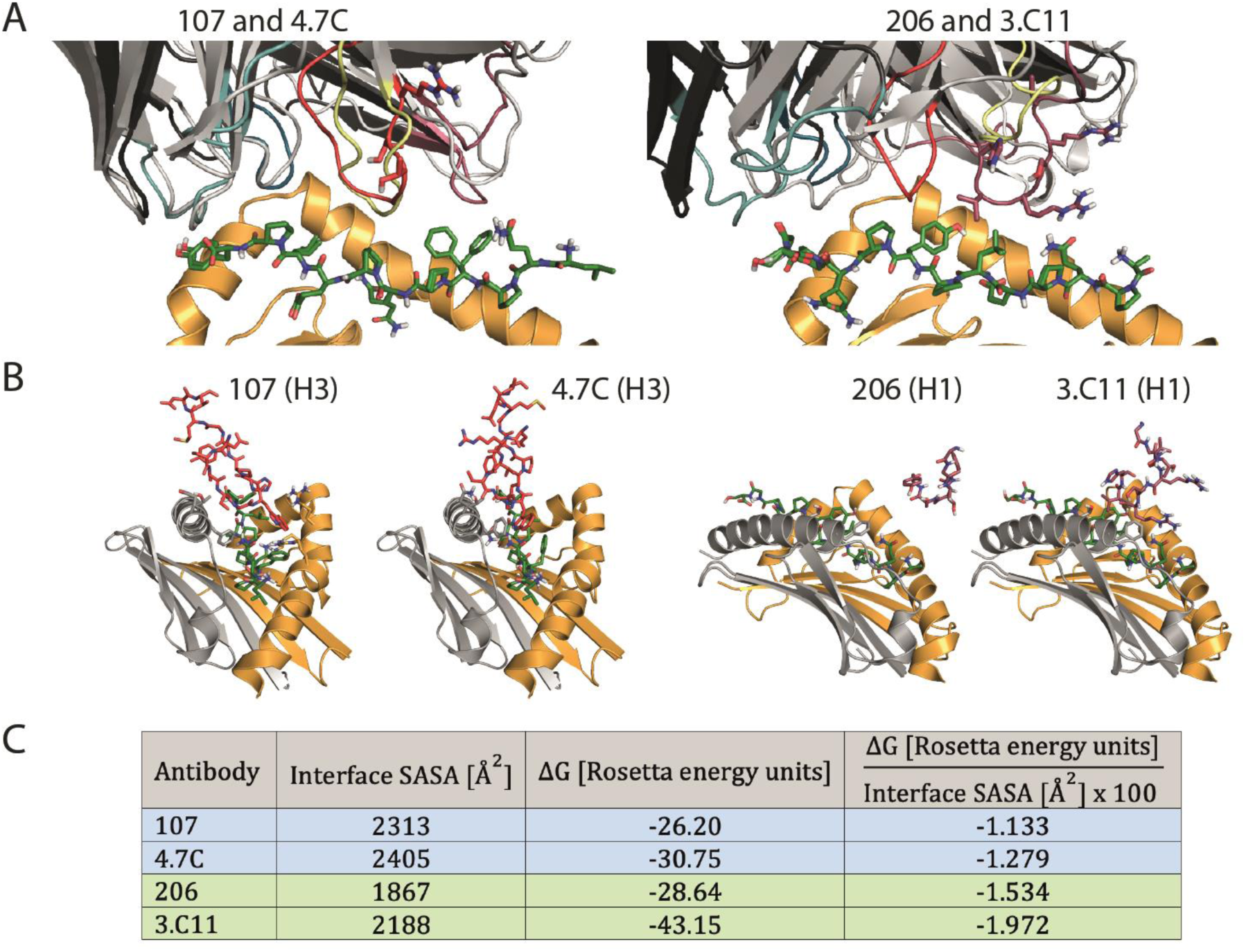
Docking models provide structural explanations for increased affinities. Models of the affinity matured variants, 4.7C and 3.C11, were generated and compared to the models of the mother clones. **A:** The high affinity variants (framework: black, light chain CDR loops: teal, heavy chain CDR loops: red) are superimposed with their respective mother clones (light grey, CDR H3: yellow). Acquired mutations are represented as sticks. **B:** The mutated CDR loops are shown as sticks in the same orientation for mother clones and high affinity variants. **C:** The solvent accessible surface area (SASA) buried at the interface and the binding energy (ΔG) were estimated from the docking models using Rosetta and the ratio of binding energy to surface area was calculated.

Interestingly, the mutations responsible for the increased affinity of 4.7C were not suggested to be directly involved in binding to the pMHC, but rather to stabilize the CDR H3 loop with additional hydrogen bonds, and the overall binding mode did not change. In contrast, 3.C11 was predicted to form several new interactions with pMHC via the CDR H1 loop as a result of the increased loop length (2 amino acids). In the model, the mutated CDR H1 loop was positioned where the CDR H3 loop was in the mother clone, which was displaced to the periphery of the interface. The interfaces were further analyzed using Rosetta’s InterfaceAnalyzer, showing the affinity matured leads produced models with consistently lower scores and larger buried surface area (**Fig. 5**C).

The solvent accessible surface area (SASA) buried at the interface increased from the mother clones to the high affinity variants in both cases, with a larger gain for the DQ2.5-glia-α2-specific antibodies. Rosetta further estimated lower binding scores (defined as the difference in score between the bound and unbound states) for the improved variants. The binding scores also improve when normalized with respect to the SASA buried at the interface, suggesting that the gains in affinity were both due to a larger interface surface and stronger interactions across the interface.

### Detection of cell-surface pMHC

Having demonstrated specificity and improved affinity of the antibodies to soluble, recombinant pMHC molecules, we tested whether the antibodies would bind pMHC complexes on the surface of cells. We loaded HLA-DQ2.5^+^ Raji B lymphoma cells (27) with soluble gliadin peptides (**Fig. 6**A). To exclude species-dependent cross-reactivity, the antibodies were reformatted and expressed as mIgG2b (S. 6A). Whereas both mother clones failed to show detectable binding to peptide-loaded Raji cells, the high affinity variants, 4.7C and 3.C11, both exhibited specific binding after loading with DQ2.5-glia-α1a or DQ2.5-glia-α2 peptides, respectively.

**Fig. 6:**
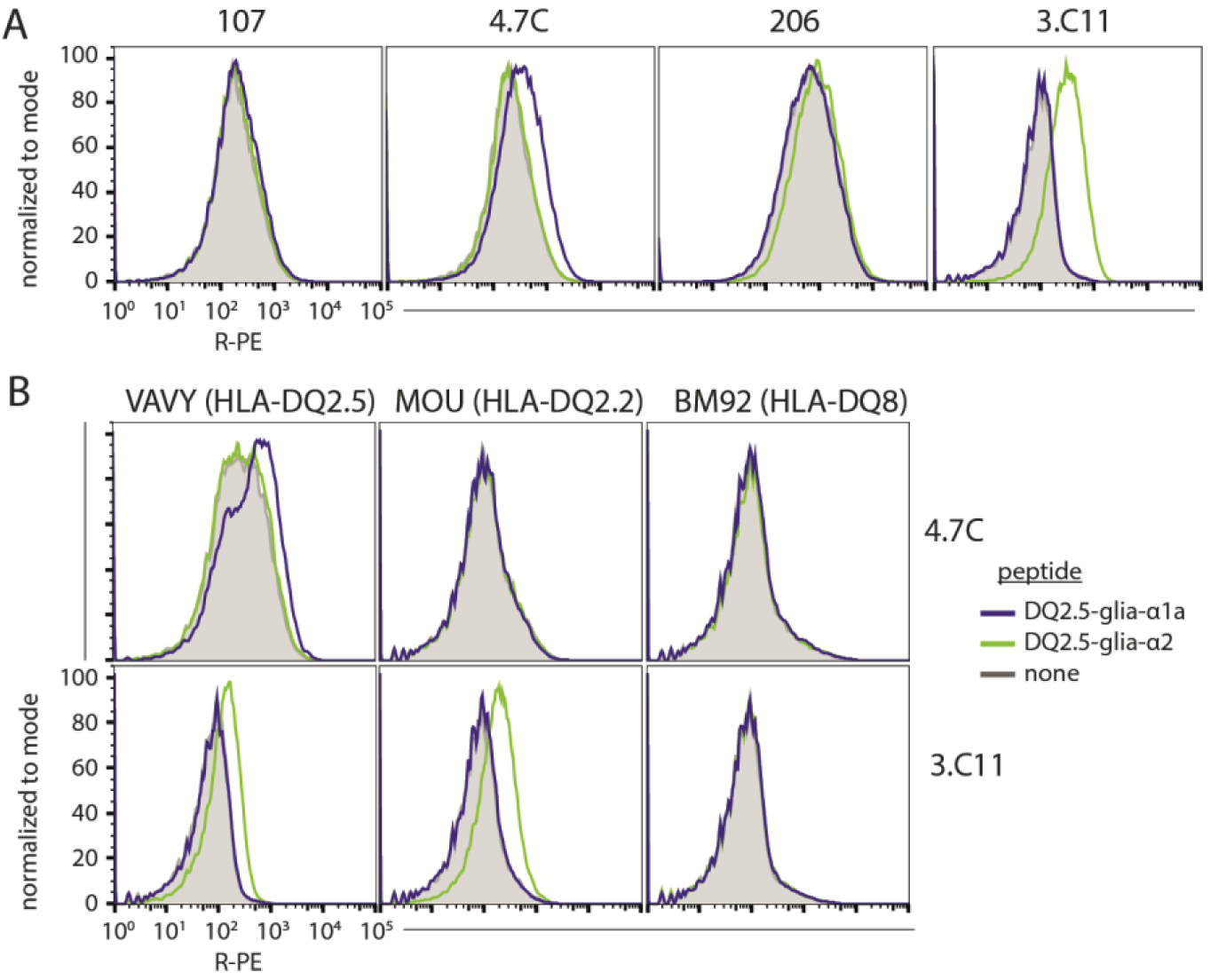
The pMHC-specific antibodies bind peptide-loaded human B-cell lines. **A**: Raji B cells were *in vitro* loaded with 50 µM gluten peptides as annotated and stained with 107, 4.7C, 206, and 3.C11 mIgG2b conjugated to R-PE**. B:** Human EBV transduced B-cell lines with different HLA-DQ allotypes were loaded with gluten peptides and stained as before (n=2).

To further assess the HLA restriction of the antibodies, human EBV transduced B-cell lines of different HLA-DQ allotypes were loaded with gluten peptides and immediately stained (**Fig. 6**B, S. 6B). HLA-DQ2.2 binds α-gliadin epitopes less stably than HLA-DQ2.5, and HLA-DQ8 is not expected to bind them at all (28). The flow cytometry experiments confirmed that the antibodies were pMHC-specific, with 4.7C being restricted to HLA-DQ2.5 as previously shown for its mother clone 107 (18), and 3.C11 binding to peptide presented on either HLA-DQ2.5 or the closely related HLA-DQ2.2. Neither of the antibodies bound HLA-DQ8.

### Staining human small intestinal biopsy material

We recently showed that the DQ2.5-glia-α1a-specific mother clone of the high affinity 4.7C antibody specifically stained B cells and PCs in single-cell suspensions made from inflamed intestinal biopsies derived from HLA-DQ2.5^+^ CeD patients who consumed gluten (18). Here, we again generated fresh single-cell suspensions from such HLA-DQ2.5^+^ CeD patients that were on a gluten containing diet (untreated), or control subjects with normal gut histology. We then stained with the mIgG2b versions of 4.7C and 3.C11, as well as antibodies with different APC surface marker specificities (**Fig. 7**). Indeed, we confirmed the previous observation using the more sensitive high-affinity antibodies. Both specificities exhibited roughly the same staining pattern and level (**Fig. 7**A+B, S. 7). The results of the staining of biopsies from three control subjects suggested that there was little background staining with all antibodies used. Two out of eight patient samples were seemingly negative for the targeted pMHC using both antibodies. We further extended the analysis to pMHC expressed on DCs and Mfs (**Fig. 7**C+D). The pMHC-specific antibodies did not stain cells in control subjects, but they appeared to detect low levels of peptide presentation in untreated HLA-DQ2.5^+^ CeD patients. Of note, the total number of DC/Mf was low in the biopsy material.

**Fig. 7:**
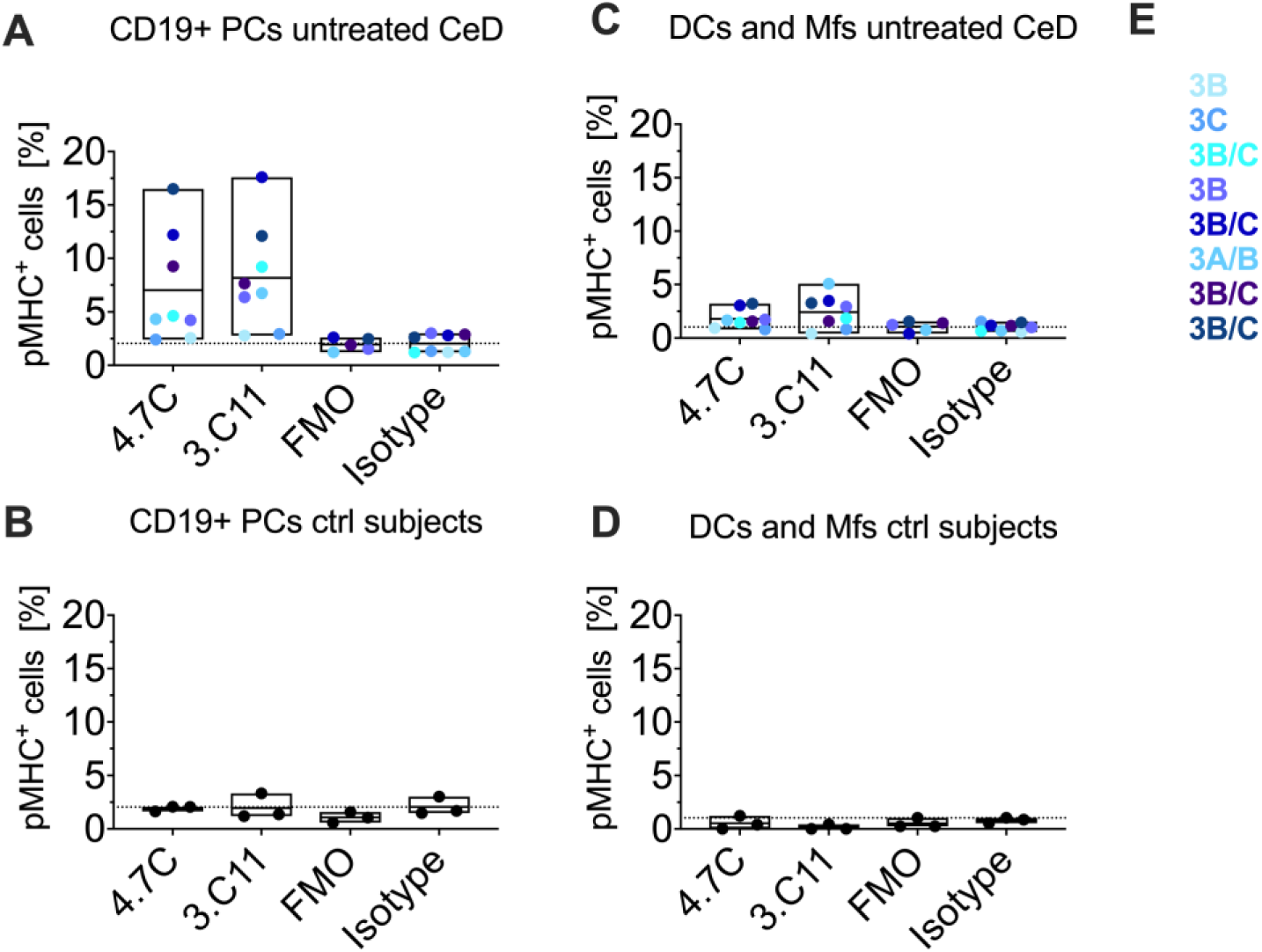
The pMHC-specific antibodies detect gluten peptide presentation on cells derived from small intestinal biopsies from CeD patients. Single-cell suspensions were prepared from either untreated HLA-DQ2.5^+^ CeD patients (n=8) (A+C) or controls with a normal intestinal histology (n=3) (B+D). Cells were gated as live, large lymphocytes, CD3^−^CD11c^−^CD14^−^CD38^+^CD27^+^CD19^+^CD45^+^ PCs (A+B) or as live, CD3^−^CD19^−^CD27^−^CD38^−^CD11c^+^CD14^+^ DCs and Mfs (C+D). Bound mIgG2b antibodies were detected with an Alexa-546-conjugated secondary antibody and the frequency of positive cells was calculated based on gates set according to the staining of an isotype control antibody (isotype). Secondary antibody only (FMO) was also included as a control. The mean percentage in each group is shown as horizontal lines and the dotted lines represent mean background staining of the isotype control. Each CeD patient is represented by a unique color. **E:** For each CeD subject (different shades of blue), alterations in biopsy histology according to modified Marsh scores are indicated.

### High affinity antibodies are potent inhibitors of T-cell activation

Having confirmed that cell-surface pMHC was specifically detected both on *in vitro* peptide-loaded cells and on patient-derived APCs loaded *in vivo*, we wondered if we could block the interaction between T cells and APCs. We focused on the DQ2.5-glia-α2 epitope, as it is the most extensively characterized gliadin T-cell epitope to which all patients mount a T-cell response (29).

To determine the inhibitory capacity of 3.C11, human SKW3 T cells devoid of endogenous TCR were engineered to express TCRs that were cloned from CeD patients, and that were specific for HLA-DQ2.5 with DQ2.5-glia-α1a (TCR 380 (30)) or DQ2.5-glia-α2 (TCR S16 (20) and TCR 364 (31)). We loaded Raji B cells with titrated amounts of stimulatory gliadin peptide and co-cultured them with the SKW3 T cells. T-cell activation was measured by determining CD69 expression on the SKW3 T cells using flow cytometry (S. 8). A peptide concentration inducing 60% T-cell activation was chosen for subsequent experiments, where the peptide-loaded Raji B cells were incubated with 3.C11 or control antibodies upon addition of SKW3 T cells (**Fig. 8**A). Indeed, 3.C11 specifically and completely inhibited T-cell activation of the SKW3 364 T cells specific for DQ2.5-glia-α2, while the DQ2.5-glia-α1a specific SKW3 380 T cells were unaffected. The effect was HLA-DQ-specific as inhibition was observed using a pan HLA-DQ antibody (SPV-L3), whereas no effect was seen with a pan HLA-DR antibody (L243). We next assessed the specificity of this inhibitory capacity by loading Raji B cells separately with DQ2.5-glia-α1a or DQ2.5-glia-α2 or a mix of both peptides. We further included the 33mer gliadin peptide, that contains three overlapping copies of the DQ2.5-glia-α2 epitope and is suspected to be present and immunogenic in the CeD lesion (32). The peptide loaded Raji B cells were co-cultured with a blend of SKW3 380 T cells and CFSE labelled SKW3 364 T cells (**Fig. 8**B). As seen, 3.C11 specifically inhibited activation of the SKW3 364 T cells but did not affect epitope specific activation of the SKW3 380 T cells, not even when the epitopes are presented on the same HLA allotype by the same cell population. To test if this effect also held true for an unrelated T-cell response restricted to a different HLA allotype, we repeated the assay using HLA-DQ2.5 and HLA-DP4 positive K562 cells, loaded with antigenic peptides, in combination with SKW3 364 T cells or the MAGE-A3 specific SKW3 R12C9 T cells restricted to HLA-DP4. As in the previous assay, 3.C11 fully blocked SKW3 364 T cells whereas there was no effect on the SKW3 R12C9 T cells.

**Fig. 8:**
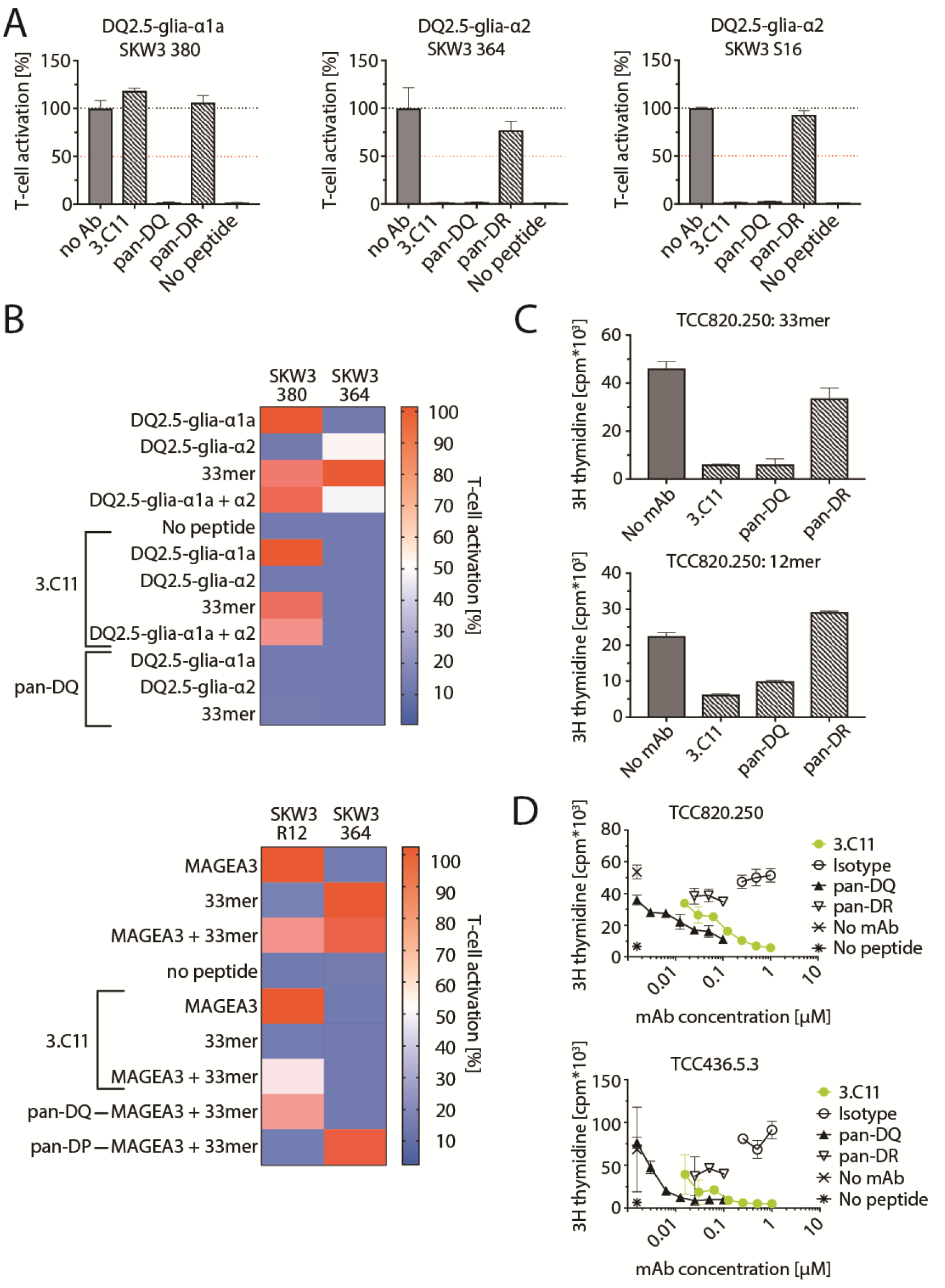
Inhibitory capacity of the HLA-DQ2.5:DQ2.5-glia-α2-specific antibody 3.C11. **A:** At a peptide dose leading to 60 % of maximum T-cell activation as measured by CD69 upregulation, 1 µM pMHC-specific antibodies or 0.1 µM pan-HLA antibodies were added to the Raji B cells prior to incubation with T cells. T-cell activation was calculated relative to peptide-specific T-cell activation without presence of antibodies. **B:** Raji B cells were loaded with gliadin peptides or combinations of them and co-cultured with a blend of SKW3 T-cells 380 and 364 (***upper panel***). K562-CIITA cells were loaded with stimulatory peptides or combinations of them and co-cultured with a blend of the two SKW3 T-cells 364 and R12C9 **(*Lower panel*)**. In both panels, SKW3 364 T cells were labelled with CFSE to guide T-cell separation in flow analysis in presence or absence of antibodies as annotated. Signals were normalized to antigen-specific activation in absence of antibodies. Error bars illustrate mean ± SD of duplicates **C:** The primary T-cell clone (TCC) specific for DQ2.5-glia-α2, TCC820.250, was co-cultured with peptide peptide-loaded Raji B cells in presence of antibodies as annotated. T-cell proliferation was assessed in a thymidine incorporation assay. **D:** HLA-DQ2.5:DQ2.5-glia-α2-specific TCCs (TCC820.250 and TCC4356.5.3) were co-cultured with 33mer-loaded Raji B cells in presence of titrated amounts of antibodies as annotated. Pan-HLA-DQ antibody (SPV-L3), pan-HLA-DR antibody (L243 or B8.11), HLA-DP antibody (B7/21), and an isotype were included as controls as annotated.

Finally, we tested the effect of 3.C11 on proliferation of primary T-cell clones (TCCs) derived from CeD patients (**Fig. 8**C). We co-cultured the DQ2.5-glia-α2-specific TCC, TCC820.250, with Raji B cells loaded with 12mer DQ2.5-glia-α2 or the 33mer gliadin peptide. In line with the inhibition of the reconstructed SKW3 T cells, we here observed a complete inhibition of T-cell proliferation for both the minimal epitope and the highly stimulatory 33mer peptide. We then performed a similar experiment using two different DQ2.5-glia-α2-specific TCCs and added titrated amounts of antibodies (**Fig. 8**D). Antibody 3.C11 inhibited T-cell proliferation in a concentration-dependent manner, while control antibodies had minimal to no effect.

## Discussion

We generated two lead antibodies specific for the two α-gliadin pMHC complexes HLA-DQ2.5:DQ2.5-glia-α1a (18) and HLA-DQ2.5:DQ2.5-glia-α2. Both were affinity matured by phage display selection from second-generation libraries using a semi-rational strategy based on molecular docking models. The high-affinity clones were target-specific and outperformed their mother clones in affinity and sensitivity for detection of cell-surface pMHC. These reagents are valuable tools for the study of antigen presentation in CeD patients. They allowed us to confirm and extend the results from previous studies that focused on one epitope only, and enabled detection of cells with low levels of peptide presentation. Furthermore, they inhibited T-cell activation *in vitro* and may thus have therapeutic potential.

The primary leads were selected from a human naïve scFv library that contains diverse light and heavy chain gene segments (19). However, the light chain sequences of the two mother clones, 107 and 206, were remarkably similar. The models predicted conserved sequence stretches in the CDR L1 and L3 loops to interact with the MHC α- and β-chains in a similar manner for both mother clones. We speculate that the light chains might be central in recognition of the MHC molecule, while the heavy chain CDR loops primarily confer peptide-specificity. Whereas the three DQ2.5-glia-α1a lead candidates for specific binding to the DQ2.5-glia-α1a complex used the germline segments IGHV6-1 and IGKV1-9 (Supplementary Table 3), the eight lead candidates against the DQ2.5-glia-α2 complex all used the VH1-69 segment and diverse IGKV segments. Thus, our data suggest a peptide-directed preference primarily for VH germlines to acquire specificity, and the fact that we employed an antibody library based on a diverse, endogenously derived gene repertoire may have been crucial for the success of the primary selections.

The binding orientations of both mother clones were predicted to be diagonal across the peptide binding groove, but skewed towards the C-terminus of the peptide and the MHC α-chain when compared to crystal structures of TCRs with the same specificity (20). This agrees with results from studies showing that TCR-like antibodies generally bind diagonally, but can have more diverse orientations than TCRs (33–36). Furthermore, it illustrates that other orientations for specific recognition of pMHC are possible than what is typically seen with naturally occuring TCRs, which may be restricted by co-receptor engagement and germline-imprinted MHC recognition motifs (37).

The structural hypotheses suggested by the models were used as the basis for a semi-rational mutagenesis strategy to generate secondary scFv phage libraries. For the 107 mother clone, we also generated a fully random library with mutations distributed over the entire Fv domain, which was selected in parallel with the CDR targeted libraries. Both approaches yielded binders with increased affinity. However, the clones isolated from the CDR targeted libraries were superior. The targeted libraries were designed to primarily introduce new interactions in loops that were predicted to interact poorly with the pMHC. By comparison, other studies have successfully chosen to randomize suboptimal side chains directly neighboring hot spot residues in the main contributing loops (35, 38, 39). We found that our strategy is likely capable of introducing new interactions and increasing the interface SASA considerably, which is not the case for strategies aiming exclusively at improving existing interactions. Our strategy exploits the principle that the interface SASA is positively correlated with affinity (40, 41).

The second-generation libraries were displayed on phage coat protein pIX and selected for high affinity variants. This selection provides the first example of affinity maturation using signal sequence independent display on phage coat protein pIX. In a previous study, we displayed scFv at HV on pIX and observed efficient enrichment of full-length functional scFvs during selection. We retrieved clones with higher affinity and thermostability in direct comparison to both LV and HV display on pIII (42), suggesting that HV display on pIX could be particularly valuable for affinity engineering.

We observed staining of patient biopsies using both 4.7C and 3.C11, and primarily detected PCs which are characteristically expanded in celiac lesions. Other APCs such as Mfs and DCs are scarce in the lamina propria and we detected only little staining of the few CD14^+^ or CD11c^+^ cells using our TCR-like antibodies. Mf and DC subsets were previously shown to occur at characteristically altered frequencies in the lamina propria of CeD patients and were therefore suggested to contribute to CeD pathogenesis (43–46). Extending the number of subjects in conjunction with e.g. cell enrichment protocols may illuminate possible differences in peptide presentation.

3.C11 specifically inhibited T-cell activation and proliferation *in vitro*. This ability may be a valuable tool for future studies of CeD pathogenesis at single epitope resolution. Antibodies specific for pMHC have previously been suggested as potential therapeutics in other autoimmune disease (9, 47, 48) and cancer (49, 50). Antibodies targeting type 1 diabetes-associated autoantigens bound to human or murine MHC class II molecules have been engineered and found to exhibit T-cell inhibitory capacity *in vitro* and in mouse models (9, 47). Decreased activation of antigen-specific T cells in lymph nodes and spleen has been observed in immunized mice treated with antibody (9). Treatment with another pMHC-specific antibody delayed disease onset and progression, which could be attributed to a decrease in lymphocytes infiltrating the pancreatic islets. This observation was disease-specific, but not restricted to insulin-specific T cells and seen for both CD4^+^ and CD8^+^ T cells, as well as B cells (47). This suggests that an antibody inhibiting a single T-cell epitope may be sufficient to induce a more tolerogenic immune response, which would be beneficial in the treatment of autoimmune diseases, including CeD. Thus, the use of high-affinity TCR-like antibodies such as the ones reported in this study may offer a therapeutic approach through single epitope targeting (29, 32)(51). Collectively, our findings encourage further studies to assess the therapeutic potential of these gliadin-specific TCR-like antibodies for treatment of CeD.

## Material and Methods

Methods for antibody engineering are described in detail in the SI Materials and Methods. Briefly, first-generation HLA-DQ2.5:DQ2.5-glia-α2-specific antibodies were isolated from a human naïve scFv library (19) selected on recombinant pMHC essentially as described before (18, 42).

Structural models of antibodies bound to the pMHC complexes were generated using RosettaAntibody and SnugDock (52). Targeted phage libraries were constructed using degenerate oligonucleotides and scFv were displayed on phage coat protein pIX (24) and selected for high affinity and stability using a protocol based on (25). Candidate clones were assessed in nanoDSF, ELISA, SPR, and flow cytometry experiments. Inhibition of T-cell activation and proliferation was assayed by reading out CD69 expression of retrovirally transduced gliadin-specific T-cells in flow cytometry and in thymdine incorporation assays using primary TCCs, respectively.

## Author Contributions

RF, LSH, IH, SK, GB, JRJ, KSG, TF, and ESV designed and performed research and analyzed data. KEAL and JJ performed endoscopic examination and provided biopsies. SY provided intestinal resections. LSH, JJG, LMS, IS, and GÅL designed research, analyzed data and supervised the study. RF, LSH, IS, and GÅL wrote the manuscript. All authors critically reviewed the manuscript.

## Acknowledgement

We would like to thank Sathiaruby Sivaganesh and Marie K. Johannesen for excellent technical assistance, Ute Krengel for helpful discussions, The Flow Cytometry Core facility at Oslo University Hospital for assistance with cell sorting and the Department of Immunology at Oslo University Hospital – Rikshospitalet for HLA-typing.

This work received funding from the South-Eastern Norway Regional Health Authority (grants 2016039 and 2018067) and the Research Council of Norway through its Centers of Excellence funding scheme, project number 179573/V40 and Stiftelsen Kristian Gerhard Jebsen. JRJ and JJG are supported by the U.S. National Institutes of Health grant R01-GM078221. The computations were performed on resources provided by the Maryland Advanced Research Computing Center (MARCC) and UNINETT Sigma2 – the National Infrastructure for High Performance Computing and Data Storage in Norway.

## Supplementary Materials and Methods

### Recombinant pMHC expression and purification

Recombinant, soluble pMHC with the peptides DQ2.5-glia-α1a (QLQ<u>PFPQPELPY</u>), DQ2.5-glia-α2 (<u>PQPELPYPQ</u>PE), CLIP2 (MAT<u>PLLMQALPM</u>GAL), DQ2.5-glia-ω1 (QQ<u>PFPQPEQPF</u>P), DQ2.5-glia-ω2 (F<u>PQPEQPFPW</u>QP), DQ2.5-glia-γ2 (QGI<u>IQPEQPAQL</u>), and DQ2.5-hor-3 (EQ<u>PIPEQPQPY</u>P) covalently linked to the MHC β-chain (linker GAGSLVPRGSGGGGS) were produced in insect cells and affinity purified using the monoclonal antibody 2.12.E11 as previously described (1). The recombinant pMHC molecules were biotinylated in a site-specific manner using the enzyme BirA. If used for SPR, monomeric proteins were isolated by size exclusion chromatography using a Superdex 200.

### Antibody modeling

Structural models of Fv fragments were generated as described (2, 3). CDR loops and framework region templates were selected by sequence homology and combined to produce a single “grafted” model using RosettaAntibody. We used 10 templates for the V_L_/V_H_ relative orientation (4) resulting in 10 grafted models. The grafted models were then energy minimized (relaxed) in the Rosetta score function (5, 6) and further improved by *de novo* CDR H3 loop modeling and V_L_/V_H_ docking. The CDR H3 loop was constrained to a kinked conformation (7) and a total of 2,800 Fv models were generated. The 10 final models were selected based on low Rosetta score and V_L_/V_H_ orientations within the natural distribution. Models were taken from at least three initial grafted templates to maintain diversity.

### Antibody docking to pMHC

We used the cocrystal structure with the T-cell receptor (4OZI) (8) as a template for an initial orientation of the Fv models relative to the pMHC. Crystal structures of HLA-DQ2.5:DQ2.5-glia-α1a (1S9V) (9) and HLA-DQ2.5:DQ2.5:glia-α2 (4OZF) (8) were retrieved from the PDB and relaxed in the Rosetta score function. Combining each HLA structure with the corresponding ensemble of 10 final Fv models, we used SnugDock to generate 1,000 complex models, as described in (10)structurally similar models separated from the bulk of models by a decrease in score), and an agreement with the experimental observations. When funnels were detected, the lowest scoring representative model was chosen to prepare figures. Rosetta’s InterfaceAnalyzer application was used to obtain information about binding scores and interfaces (11–13).

### Primary phage display selection

The HLA-DQ2.5:DQ2.5-glia-α2-specific **mother clone 206** was isolated from a naïve fully human scFv library reformatted to the pFKPDN vector (14) for display on coat protein pIII along with co-expression of the chaperone FkpA (15). The selection protocol was essentially as described in (3, 15). Phage input in R1 was 1.0×10^11^. A total of three rounds of selection were performed. Phage libraries, beads and tubes were blocked with 5% skim milk or 2 % BSA in PBS supplemented with 0.05% tween-20 (PBS-T) prior to each round of selection. Blocked phages were incubated with biotinylated HLA-DQ2.5:CLIP2 (negative selection) in volumes of 1 mL PBS-T in 1.5 mL tubes on a rotating wheel for 1 h at room temperature and removed bound phages with strepavidin-coated magnetic beads (Dynabeads MyOne Streptavidin T1, Invitrogen). Next, phage libraries were selected on biotinylated HLA-DQ2.5:DQ2.5-glia-α2 with decreasing concentrations of 160 nM, 16 nM and 1.6 nM from R1 to R3 before immobilization on magnetic beads. Bound phage particles where washed with PBS-T and PBS (5+5 washes in R1, 10+10 washes in R2 and 20+20 washes in R3 of selection). Phages were eluted by incubation in 500 µL 0.5% trypsin (Gibco) and incubation at room temperature on a rotating wheel for 10 min. Half of the eluted phage were used for infection of 50 mL cultures of fresh log-phase *E. coli* SS320 in 2x YT medium supplemented with 30 µg/mL tetracycline, 0.15 M glucose (2x YT-TG) at 37 °C/80 rpm/30 min. *E. coli* were pelleted, resuspended in 1 mL 2x YT and plated onto LB-agar Bio-Assay dishes (245 mm × 245 mm, Nunc) supplemented with 30 µg/mL tetracycline, 100 µg/mL ampicillin, and 0.1 M glucose (LB-TAG) as above and grown at 30 °C/16 h. *E. coli* were scraped and reinoculated to OD 0.05 in 2x YT-TAG and grown at 37 °C/200 rpm until OD reached 0.1-0.2. Cultures were superinfected with helper phages M13KO7 for low valence display or Hyperphage for high valence display at MOI 10. Cells were incubated 37 °C/30 min/80 rpm followed by 37 °C/30 min/200 rpm. Medium was then changed to 2x YT supplemented with 100 µg/mL ampicillin and 50 µg/mL kanamycin (2x YT-AK) and phage were packaged at 37 °C/200 rpm/7 h. Cells were pelleted, and supernatants filtered (0.22 µm) and PEG/NaCl precipitated overnight (ON), before resuspension in PBS.

### Screening of primary selection output

Screening of selection output was done after batch cloning the selection output into pFKPEN (14) for soluble scFv expression as indicated. 96 clones from the R3 output were picked into a 96-deep-well plate containing 400 µL 1x LB-AG and incubated at 37 °C ON shaking. 50 µL of ON cultures were transferred to a fresh 96-deep-well plate containing 400 µL 1x LB-AG and incubated 4 h/37 °C/600 rpm. Cells were harvested by centrifugation and resuspended in 450 µL 2x YT-A supplemented with 0.1 mM IPTG. Plates were incubated at 30 °C/ON/600 rpm. Cells were harvested and supernatant samples were taken. Periplasmic fractions were extracted on ice/1 h using 300 µL of 50 mM Tris-HCl, 20 % sucrose, 1 mM EDTA, pH 8, 1 mg/mL lysozyme, and 0.1 mg/mL RNaseA. Cells were pelleted and periplasmic samples taken and blended 1:1 with supernatant samples for use in ELISA.

### Generation of targeted and randomized scFv phage libraries

The **targeted libraries** were based on scFv sequences of mother clones 107 and 206 in pGALD9ΔLFN (16). A modified version of a previously described protocol was employed to generate the targeted libraries (17), now including a preparative step to remove template background, and a lower concentration of DNA in transformation to minimize multiple genotypes per cell during initial library packaging. Briefly, AgeI restriction enzyme sites were introduced into the target sequences using mutagenic oligonucleotides and a standard PCR protocol. Mother clone phage containing the AgeI sites were packaged from *E. coli* XL1-Blue by superinfection with M13K07 (MOI 10) and precipitated with PEG/NaCl (20 % w/v PEG 8,000, 2.5 M NaCl). Single-stranded (ss) DNA was isolated from phage suspensions using a QIAprep Spin M13 kit and verified with agarose gel electrophoresis. 8 sets of mutagenic oligonucleotides with 6-7 degenerate NNK codons (N: A/C/G/T, K: T/G) were designed so that they target relevant stretches of the CDR H1 or H3 loops for randomization and length variation. The oligonucleotides were 5’-phosphorylated and annealed to the ssDNA. Heteroduplex DNA was synthesized using T7 DNA polymerase and ligated with T4 DNA ligase. Covalently closed circular double-stranded DNA was purified by agarose gel electrophoresis and transformed into electrocompetent *E. coli* AVB100FmkII (18) using a BTX ECM 600. Transformants were plated onto LB-TAG agar Bioassay dishes (245 mm × 245 mm, Nunc) and grown at 30 °C/16 h. Primary transformations were spot titrated onto nitrocellulose membranes. Plates were scraped and DNA was isolated and digested with AgeI to remove template. Undigested library DNA was purified from agarose gels and transformed into electrocompetent *E. coli* SS320 (Lucigen) and again plated onto LB-TAG agar Bioassay dishes as before. Phage were packaged as described (15) using Deltaphage (19) for high valence display on coat protein pIX and precipitated with PEG/NaCl twice before R1 of selection.

The **random library** based on mother clone 107 was generated using the JBS dNTP-Mutagenesis Kit (Jena Bioscience) according to manufacturer instructions. Template DNA was removed by DpnI digest. Sequence analysis revealed that the amino acid mutation frequency was 4% and mutations were distributed uniformly along the scFv cassette. The total diversity of this library was estimated to 5×10^7^ by spot titration onto nitrocellulose membranes. Mutated scFv were subcloned into the pGALD9ΔLFN vector. DNA was purified by ethanol precipitation (Pellet Paint TM, Novagen) and transformed into *E. coli* SS320 (Lucigen) and packaged as described above.

### Phage display selection for affinity maturation

The selection of the second-generation libraries was performed similarly as described above for the primary selections. Phage input in R1 was 3.0×10^11^. The target antigen concentrations were 100 nM, 1 nM, and 10 nM in R1 to R3. Washing was increased from 5+5 in R1 to 10+10 in R2 and R3. In the thermal branch, phage were heated for 15 min to 60 °C and 65 °C in R2 and R3, respectively. After the heat challenge, phage samples were cooled down on ice and centrifuged at 17,000xg/5 min. Phage samples were transferred to new tubes for selection. In the competition branch, bead-bound positively selected phage particles were incubated with 100 nM of scFv or 50 nM of hIgG1 of the respective mother clones. All libraries were displayed on coat protein pIX and selected phages were rescued by superinfection with M13KO7 for low valence display and Deltaphage for high valence display.

### Screening of affinity matured selection output

Screening of selection output was done in soluble scFv format as described for screening of the primary selection output. For screening in phage format, single clones from the R3 output were picked into 96-deep-well plates containing 2x YT-AG and cultured ON/37 °C/600 rpm. 10 µl were transferred to fresh plates containing the same medium and grown for 3 h/37 °C/600 rpm. *E. coli* were infected with 10^9^ cfu Deltaphage per well and incubated at 37 °C/30 min with gentle shaking, and 30 min with vigorous shaking. Cells were pelleted and resuspended in 50 µl 2x YT-AK and phage were packaged ON/30 °C/600 rpm. Cells were harvested and 100 µL supernatant were used for screening in ELISA.

### Prokaryotic protein expression

Selected variable regions were either individually subcloned or batch-cloned as NcoI/Not I fragments from phagemids into the pFKPEN vector (20), that contains a His-tag as well as a Myc-tag. DNA was transformed into electrocompetent *E. coli* XL1-Blue. For protein expression, the *E. coli* were inoculated into 10 mL 2x YT-AG and incubated 37 °/ON/200 rpm. Cultures were reinoculated into 1 L 2x YT-AG to OD600nm 0.025 and grown to OD600nm 0.6. Medium was changed to 1 L 2x YT-A supplemented with 0.1 mM IPTG and incubated 30 °C/ON/200 rpm. Cells were pelleted and periplasmic fractions extracted using 80 mL periplasmic extraction solution as described in screening of primary selection output. Cells were pelleted, and periplasmic fractions were filtered, diluted by adding 100 mL PBS supplemented with 150 mM NaCl and 0.05% sodium azide, and adjusted to pH 7.4 before purification.

### Eukaryotic protein expression

Variable regions were cloned as BsmI/BsiWI fragments into pLNOH2 oriP and pLNOκ oriP genomic expression vectors encoding constant human γ1 and constant human κ domains (21). The mIgG2b variants were obtained by cloning variable regions together with mouse γ2b or κ cDNA as BsmI/BamHI fragments into the same vector. DNA was transformed into electrocompetent *E. coli* XL1-Blue and purified. HEK293E cells (ATCC) were transfected at 90% confluency with equal amounts of light and heavy chain DNA using Lipofectamine 2000 (Invitrogen) (21). Cells were cultured in RPMI supplemented with 10 % fetal calf serum (FCS), 0.1 mg/ml streptomycin and 100 U/ml penicillin. Supernatant was harvested every 2–3 days for 2 weeks. Supernatants were filtered (0.22 µm) before purification.

### Purification of antibodies and antibody fragments

Antibodies, Fab fragments, or scFvs were captured on protein L columns (HiTrap, GE Healthcare) or CH1 capture select columns (Thermo Fisher Scientific), eluted with 0.1 M glycine-HCl pH 3 and neutralized with 1 M Tris-HCl pH 8. Alternatively, scFvs were purified by IMAC (HiTrap, GE Healthcare) and eluted with 50 mM Tris-HCl, 0.5 M NaCl 0.5 M imidazol pH 7.4. Proteins were size excluded on a Superdex 200 column using PBS supplemented with 150 mM NaCl (GE Healthcare).

### ELISA

EIA/RIA plates were coated with 10 µg/ml NeutrAvidin in PBS (100 µl/well, Thermo Scientific) and incubated ON/4 °C. Plates were blocked with 5% skim milk powder in PBS-T (300 µl/well) for 1 h/RT with agitation. Equal amounts of biotinylated pMHC (normalized to 300 ng/ml) were captured for 1 h/RT and followed by addition of 0.5 µg/ml purified pMHC-specific antibodies, 5 µg/ml purified Fab fragments or purified scFvs, and varying concentrations of soluble scFv or phage during screening. Bound scFv were detected with mouse anti-Myc tag antibody (Invitrogen, 1:5,000) and anti-mouse-HRP (GE Healthcare, 1:2,000). Bound hIgG1 were detected with anti-hIgG-ALP (Sigma Aldrich, 1:3,000), Fab fragments with anti-hCκ-ALP (Sigma, 1:3,000), phage particles with anti-M13-HRP (Amersham Biosciences, 1:5,000). All antibodies were diluted in PBS-T. Plates were developed with TMB solution (Calbiochem) and read at 450 nm using a microplate reader (Tecan sunrise) after the enzymatic reaction had been stopped by addition of 1 M HCl. Alternatively, the plates were developed with 1 mg/ml phosphatase substrate (Sigma Aldrich) in diethanolamine buffer and read at 405 nm.

### Thermostability analysis by nanoDSF

Melting temperatures were assessed using a Prometheus nanoDSF (NanoTemper). 10 µL of 0.2 mg/mL purified Fab fragments in PBS were transferred to glass capillaries (NanoTemper) in triplicates. A 1 °C/min temperature gradient was applied to the samples from 20 °C to 95 °C. Excitation wavelength was 295 nm and emission was measured at 330 nm and 350 nm. Data was collected and analyzed using AB-Protein PR.ThermControl V2.12.

### Binding analysis by SPR

Kinetics of antibody binding to pMHC were determined using a Biacore T200 (GE Healthcare). Briefly, NeutrAvidin (10 µg/mL in 10 mM sodium acetate, pH 4.5) was coupled onto a CM3 sensor chip to 1000 response units (RU) by amine coupling. Soluble, recombinant, biotinylated pMHC (1 µg/mL) was then captured at approximately 80-90 RU by passing over the flow cells at 10 µL/min. Antibody samples (scFv or Fab fragments) diluted in PBS supplemented with 0.05% (v/v) surfactant P20 were run over the surface at various concentrations using either single cycle kinetics or a multi cycle method. For off-rate ranking, all samples were used at 0.5 μM. Binding experiments were performed at 25°C with a flow-rate of 30 µL/min. The surface was regenerated using either glycine-HCl pH 2.2 or diethylamine pH 11 when necessary. Binding data were buffer subtracted and NeutrAvidin-reference-cell subtracted using the T200 Evaluation Software v1.0. Kinetic constants were determined by fitting the data to a 1:1 Langmuir binding model.

### B-cell lines

The human B-cell lymphoma Raji cells express HLA-DQ2.5. The human EBV-transduced B-cell line VAVY is homozygous for HLA-DQ2.5, while MOU and BM92 are homozygous for HLA-DQ2.2 and HLA-DQ8, respectively. All cells were cultured under standard conditions in RPMI 1650 supplemented with 10% FCS, 0.1 mg/ml Streptomycin and 100 U/ml Penicillin.

### Flow cytometric analyses of stained B cells

Raji and EBV-B cells were loaded with peptide by culturing in presence of 50 µM peptide ON. The following peptides were used (9mer core epitope sequences are underlined): 33mer α-gliadin LQLQ<u>PFPQPELPYPQPELPYPQPELPYPQ</u>PQPF, 12mer DQ2.5-glia-α1a QLQ<u>PFPQPELPY</u> and either an 11mer DQ2.5-glia-α2 <u>PQPELPYPQ</u>PE or 12mer DQ2.5-glia-α2 <u>PQPELPYPQ</u>PQL. The cell lines were stained with pMHC-specific mIgG2b antibodies (5 μg/ml) directly conjugated to PE (Abcam) in the presence of human FcR blocking reagent (Miltenyi, 1:50). All stainings were performed on ice using V-bottom shaped 96-well plates and an equal number of cells were used in each staining (at least 100,000). PBS supplemented with 2% FCS was used to wash cells and for dilution of antibodies and streptavidin. Data was acquired using an Attune NxT flow cytometer and analyzed using FlowJo software v10.4.1.

### Human material

Research protocols and use of human duodenal biopsy material have been approved by the Regional Ethics Committee of South-Eastern Norway (REK Sør-Øst approval 2010/2720). All subjects gave informed written consent to donate biological material to a REK Sør-Øst approved biobank (number 2012/341). CeD diagnosis was based on the guidelines from the British Society for Gastroenterology including clinical history, anti-TG2 serological testing, HLA typing and histological analysis of small intestinal biopsies obtained by esophagogastroduodenoscopy and forceps sampling from the duodenum (22). Small intestinal resections (duodenum-proximal jejunum tissue) were obtained from non-pathological small intestine during Whipple procedure (pancreatoduodenectomy) of pancreatic cancer patients who also donated material to the biobank (number 2012/341) and gave informed, written consent. Normal histology of the duodenal mucosa was confirmed in these samples.

### Isolation of single-cell suspensions from duodenal biopsies and small intestinal resections and flow cytometry

Single-cell suspensions from duodenal biopsies or small intestinal resection were prepared as described (23) and blocked with FcR Blocking Reagent (Miltenyi Biotec) and stained as detailed in Supplementary Table 4. Propidium iodide was used for exclusion of dead cells and samples were immediately acquired on LSR Fortessa cytometer (BD).

### Retroviral transduction of human SKW3 T cells and K562 cells, and flow cytometry

TCR sequences encoding the variable domains of TCRs 380 (24), 364 (25), S16 (8), and R12C9 (26) were reconstructed by gene synthesis as human/mouse chimeric TCRs as described (24) and cloned into pMSCV (Clontech Laboratories). Retroviral transduction of the SKW3 human T cells (CLS Cell Lines Service GmbH) was performed using the Retro-X Universal Packaging System (Clontech) according to the manufacturer’s instructions. Stable, homogenous TCR-expressing SKW3 T cells were obtained by standard cell expansion and FACS sorting using a FACSAria II cytometer (BD Biosciences) based on their TCR expression levels assessed by H57-Alexa647 (Thermo Fisher Scientific) antibody staining. The TCR transduced SKW3 cells were validated for peptide-specific activation using a panel of known agonistic and antagonistic peptides, essentially as described (24). T-cell activation was measured by CD69 up-regulation assessed by anti-human CD69-APC (BD Biosciences) antibody staining. Data was acquired on a BD Accuri C6 cytometer (BD Biosciences) and analyzed using FlowJo software V10 (Tree Star). The human myeloid leukemia cell line K562 (ATCC-CCL-243) was transduced with the hCIITA::eGFP dual ORF encoding retroviral vector pMMLV (VectorBuilder GmbH) to induce endogenous HLA class II expression. Stable, homogenous HLA class II positive cells were obtained by standard cell expansion and FACS sorting using a FACSAria II cytometer (BD Biosciences) based on their dual eGFP and HLA expression levels assessed by pan-HLA class II (clone CR3/43-Alexa647) antibody staining.

### T-cell activation and inhibition assays

For T-cell activation assays 50,000 Raji B cells were incubated in RPMI/10% FCS at 37°C/ON with titrated amounts of peptide, followed by washing to remove remaining free peptide and addition of 40,000 SKW3 T cells. Cells were cultured at 37°C/ON before they were analyzed in flow cytometry. The following peptides were used (9mer core epitopes are underlined): DQ2.5-glia-α1a (QLQ<u>PFPQPELPY</u>), and DQ2.5-glia-α2 (<u>PQPELPYPQ</u>PE). As a control, Cell Stimulation Cocktail containing PMA and ionomycin (eBioscience, 1:500) was added to wells containing SKW3 T cells only. Based on the established dose-response in T-cell activation, a peptide concentration estimated to result in about 60% T-cell activation (measured as CD69 upregulation on the CD19^neg^ population) was chosen for the inhibitory assays. Following ON incubation with peptide as above and washing, 1 μM (final concentration) of either 4.7C or 3.C11 were added to the Raji cells, before T cells were added and incubation continued ON. The resulting T-cell activation was measured as above. As control Abs, either 0.1 μM (final concentration) of pan-anti-DR (clone L243: Thermo Scientific) or pan-anti-DQ (clone SPV-L3: Diatech) were added in parallel.The TCCs reactive to the DQ2.5-glia-α2 epitope were established from intestinal biopsies of CeD patients (CD436 and CD820) and have been described before. Briefly, TCC820.250 was generated by direct cloning from intestinal biopsies (27), while TCC436.5.3 was generated by limiting dilution of a T-cell line established from antigen-challenged intestinal biopsy specimens (25, 28). For T-cell activation assays Raji B cells in RPMI supplemented with 10% heat-inactivated, pooled human serum, 0.1 mM 2-Mercaptoethanol, and antibiotics were irradiated with 75 Gy before 75,000 cells/well were incubated overnight with titrated amounts of peptide at 37°C. The plates were centrifuged to remove remaining free peptide before addition of 50,000 T cells (TCC436.5.3 and TCC820.250). 3H-thymidine (1 μCi/well) was added 2 days after addition of T cells and cells were harvested after an additional 16-20 hours using an automated harvester. 3H-thymidine incorporation was measured by liquid scintillation counting. For T-cell inhibition experiments, Raji B cells were incubated overnight with 1 µM DQ2.5-glia-α2 (<u>PQPELPYPQ</u>PE) or 1 µM 33mer (final concentrations) before medium exchange and addition of titrated amounts of mAbs (from 1 µM for mAbs 3.C11 and 4.7C and from 0.1 µM for mAbs SPV-L3 (Diatech) and the pan-DR antibody B8-11 (Diatech). Methods for antibody engineering are described in detail in the SI Materials and Methods. Briefly, first-generation HLA-DQ2.5:DQ2.5-glia-α2-specific antibodies were isolated from a human naïve scFv library selected on recombinant pMHC essentially as described before.

## Supplementary Information

**S. 1:**
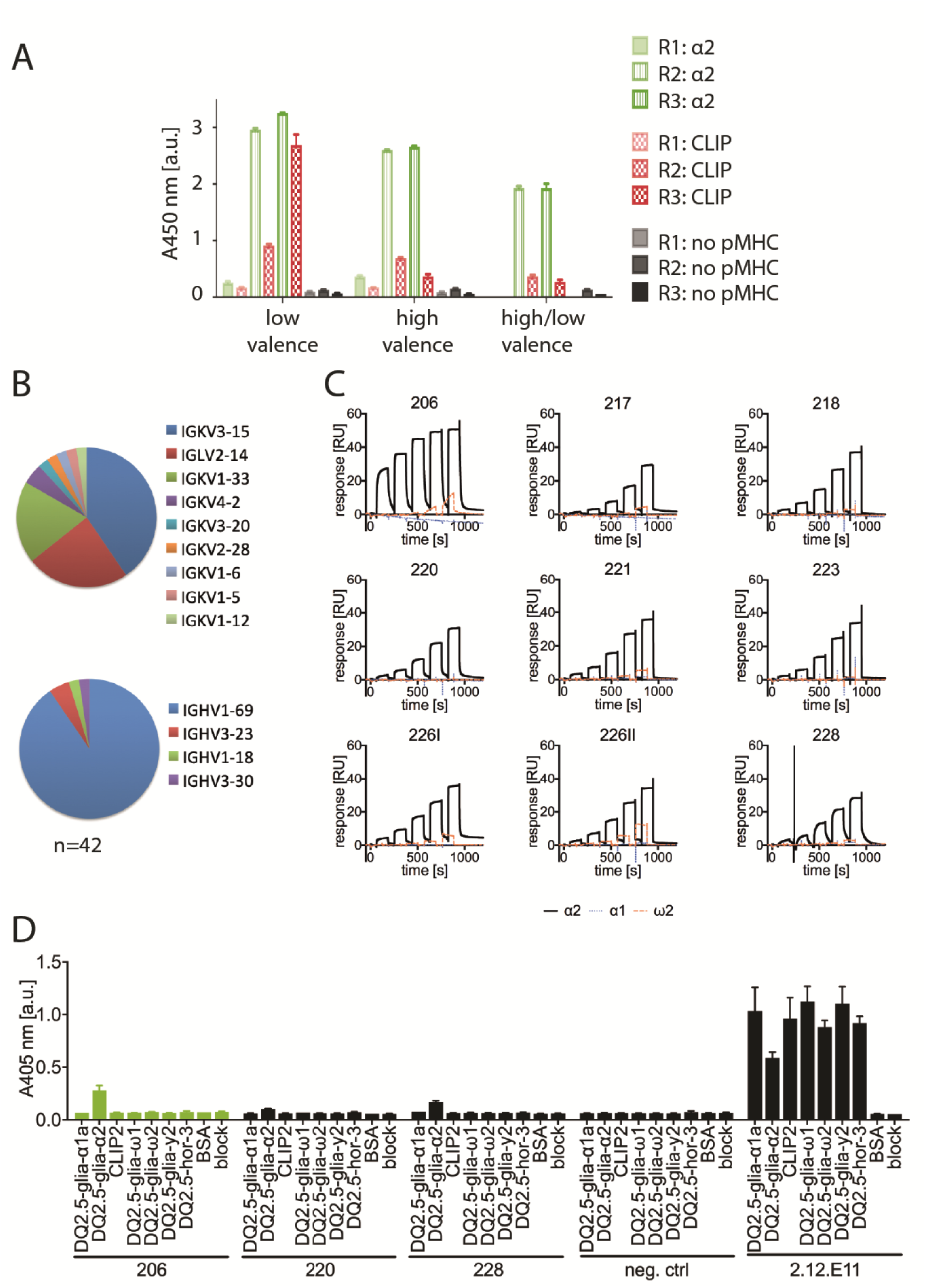
Primary selection against HLA-DQ2.5:DQ2.5-glia-a2. **A:** Polyclonal phage ELISA to assess the antigen reactivity of the phage outputs after R1-R3. HLA-DQ2.5:CLIP2, which was used for negative selection, and block was used to monitor HLA-DQ2.5 binding irrespective of peptide and background, respectively. **B:** 42 sequences of positive clones were identified after three rounds of selection and V gene usage was analyzed using IMGT/V-quest. **C:** Single scFv clones were produced in *E. coli*, affinity purified, size excluded, and analyzed in by SPR as indicated. **D:** Lead candidates 206, 220, and 228 were expressed as hIgG1 in HEK293E cells, purified on protein L columns and size excluded and then tested in ELISA for binding to different peptide MHC complexes. Functional pMHC was normalized using pan-HLA-DQ antibody 2.12.E11.

**S. 2:**
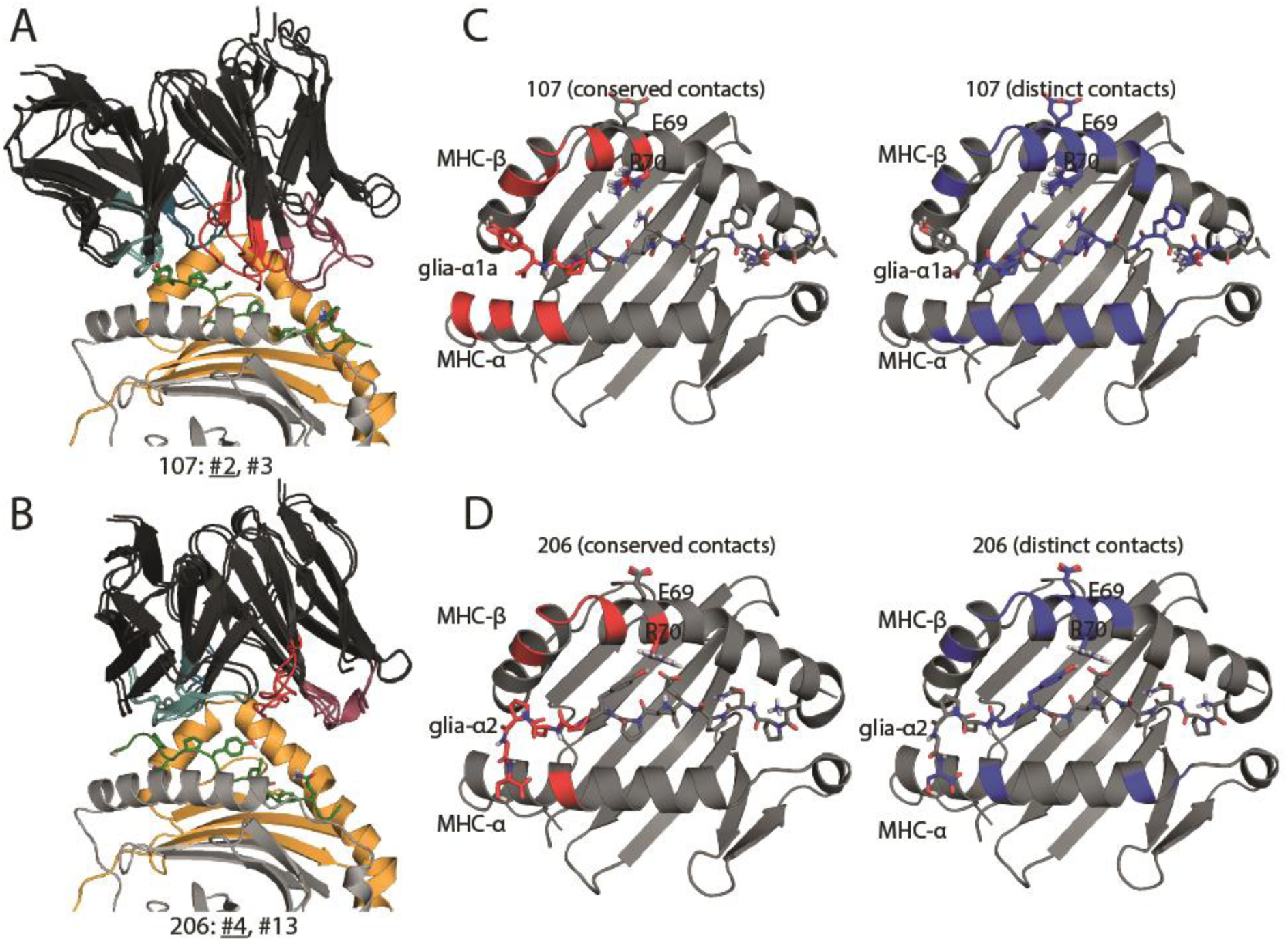
Docking models of pMHC-specific antibodies: **A+B:** 1,000 docking models were generated for 107 and 206 using SnugDock and ranked based on low Rosetta interface energy. Highly similar low energy models were manually selected and superimposed. Interface score ranks for selected models are listed and the lowest energy representative (underlined) was selected for analysis and generating figures (A+B). **C+D:** pMHC molecules are depicted as cartoons, with the peptide and crucial TCR contacts (E69 and R70) represented as sticks. The pMHC is shown in grey and the residues predicted to be within 4 Å of the antibodies are colored. Red indicates proximity to residues conserved across the two specificities and blue indicates proximity to distinct residues in the antibody variable domains.

**S. 3:**
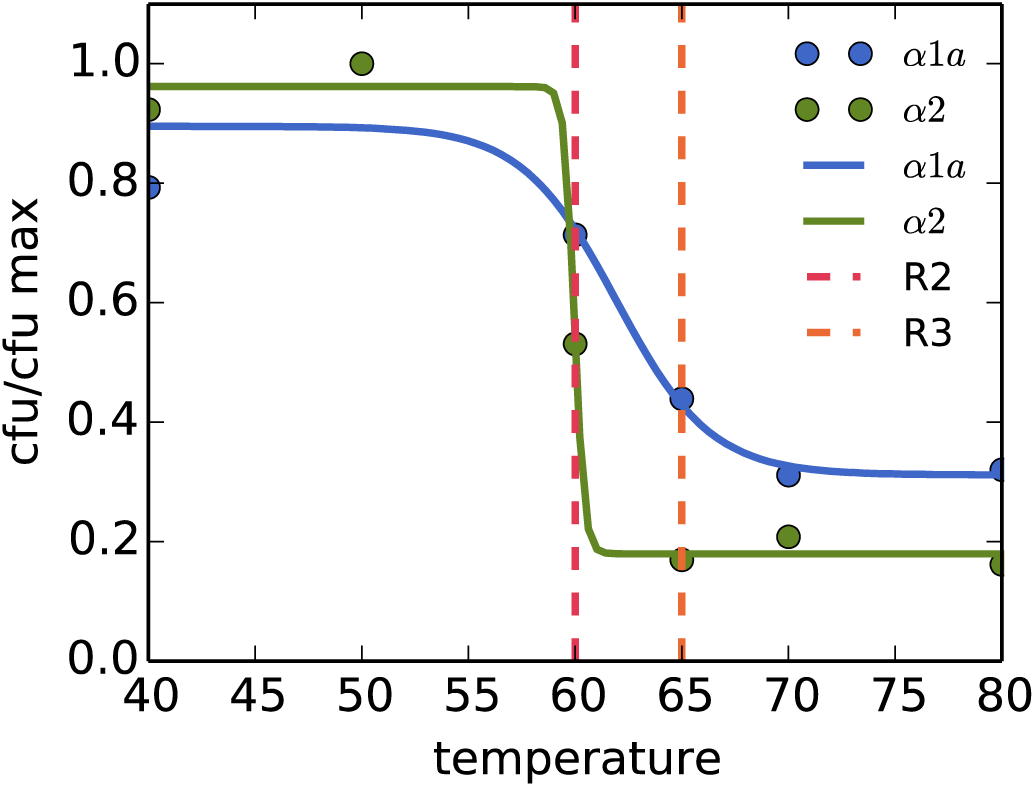
Thermostability of first-generation antibodies. The first-generation antibodies 107 and 206 were displayed at HV on phage coat protein pIX and heated at different temperatures for 10 min as annotated. They were cooled down and assessed for residual binding to protein L in ELISA and infectious titration of bound particles. We estimated melting temperatures by fitting a sigmoid function to the data (lines) and obtained 60 °C for 206 and 62 °C for 107. Heat challenge temperatures for rounds 2 and 3 are indicated for comparison (red and orange dotted lines).

**S. 4:**
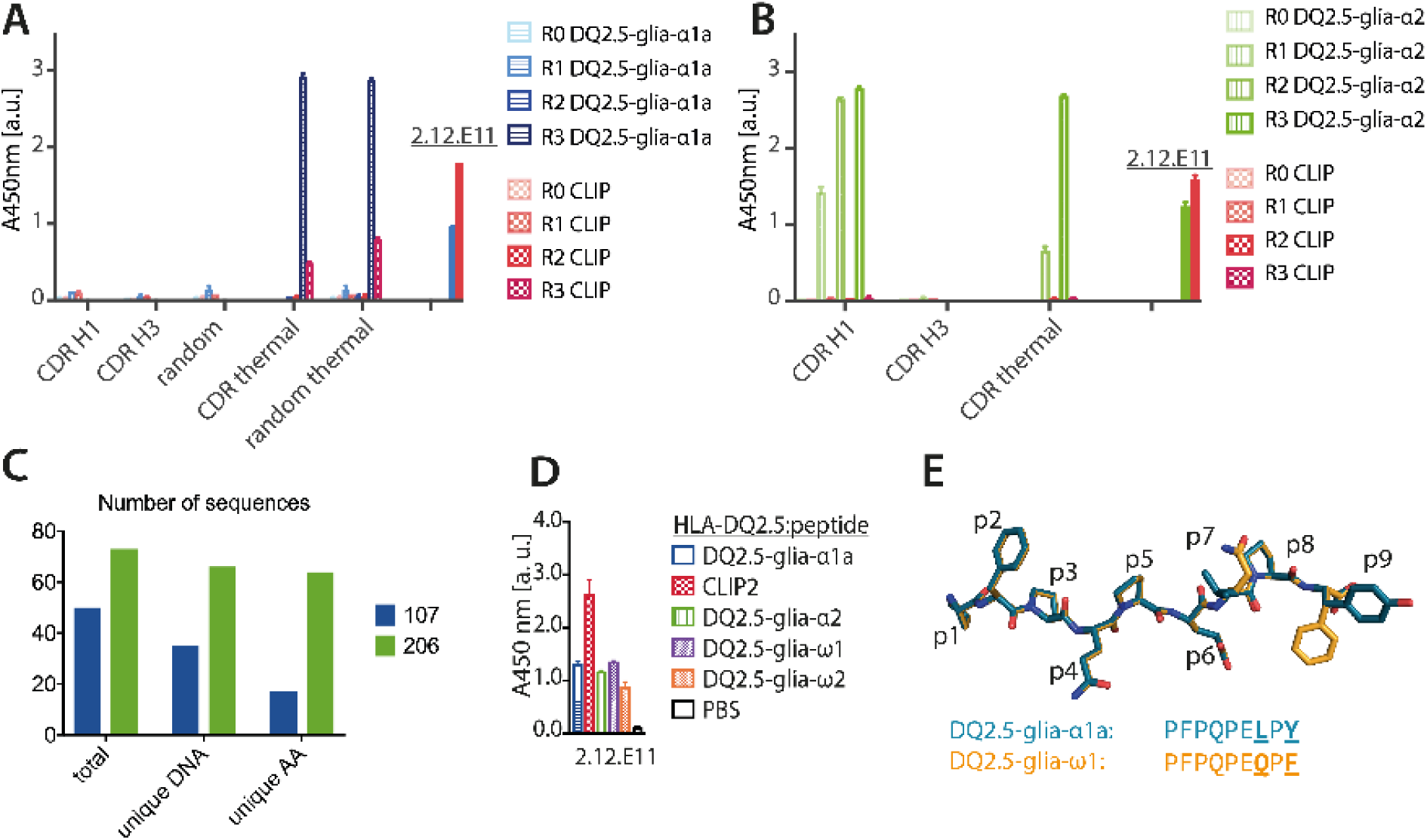
Selection for high affinity pMHC binders. **A+B:** After three rounds of selection, a polyclonal ELISA was performed to assess enrichment of binders against HLA-DQ2.5:DQ2.5-glia-α1a (A) and HLA-DQ2.5:DQ2.5-glia-α2 (B). **C:** Positive clones were sequenced and numbers of unique DNA and amino acid sequences are represented as bars. **D:** presence of functional pMHC molecules was verified by detection with 2.12.E11 in ELISA. **E:** structural alignment of DQ2.5-glia-α1a (1S9V) and DQ2.5-glia-ω1 (model). Differing positions p7 and p9 are underlined.

**S. 5:**
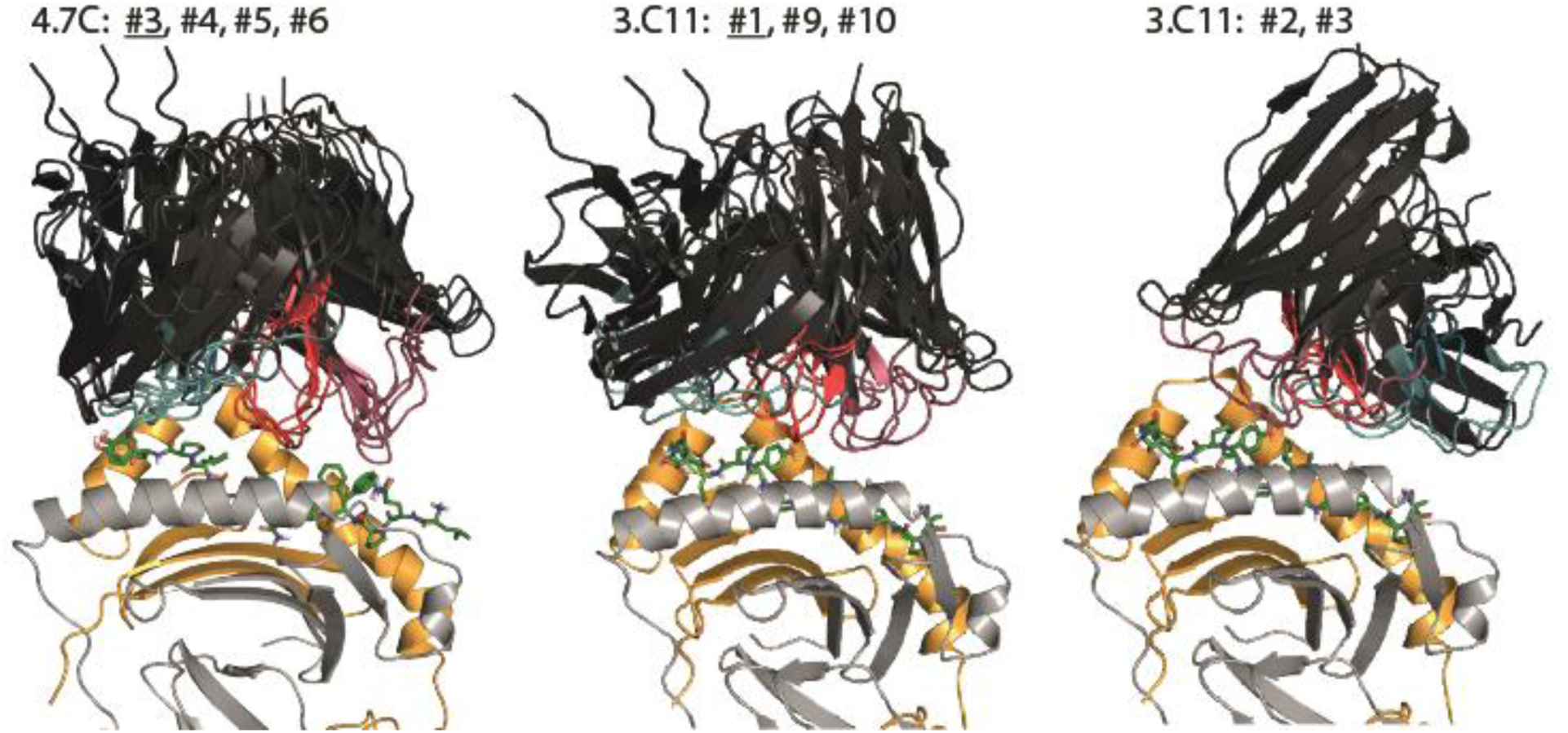
Docking models high affinity antibodies: 1, 000 docking models were generated for 4.7C and 3.C11 using SnugDock and ranked based on low Rosetta interface energy. Highly similar low energy models were manually selected and superimposed. Interface score ranks for selected models are listed and the lowest energy representative (underlined) was selected for analysis and generating figures.

**S. 6:**
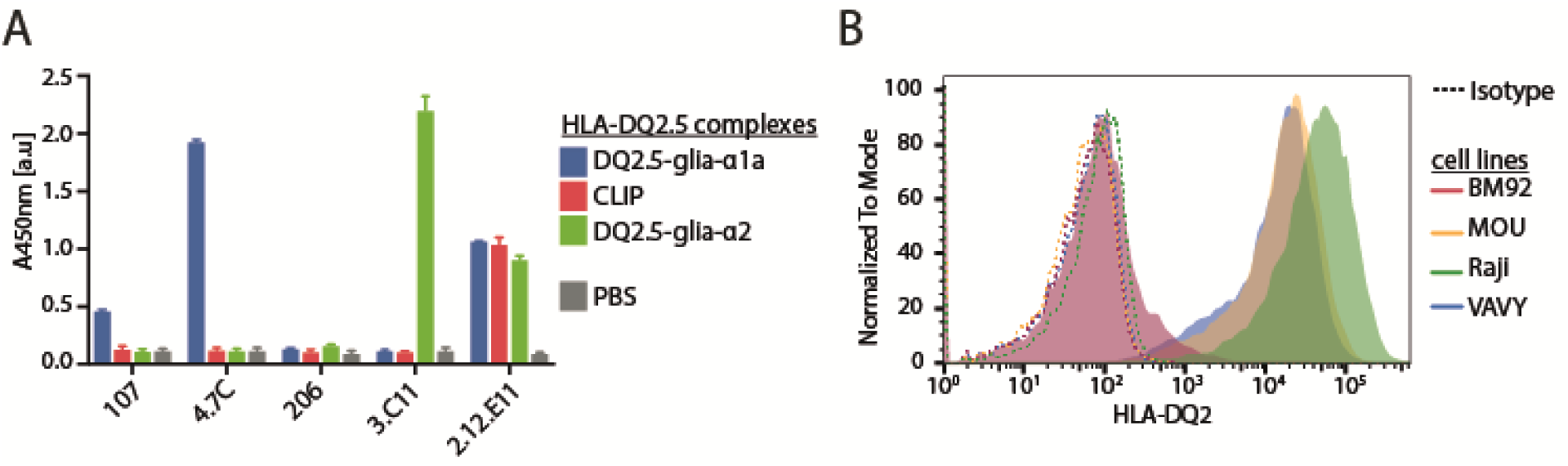
Staining for HLA-DQ2 expression. **A:** The engineered A20 mouse B cells were stained with 5 µg/mL biotinylated 2.12.E11 antibody followed by 2 µg/mL SA-R-PE. Staining with an isotype control antibody was included (dotted line). **B:** The antibodies were expressed as mIgG2b antibodies and analyzed in ELISA for binding to soluble recombinant pMHC molecules as before (0.5 µg/mL pMHC specific antibodies). The HLA-DQ2 antibody 2.12.E11 was used to control for pMHC immobilization levels. **C:** Raji lymphoma B cells and VAVY, MOU, and BM92 EBV EBV-B cells were stained with 5 µg/mL 2.12.E11 directly conjugated to R-PE (filled histograms) or an isotype control (dotted lines).

**S. 7:**
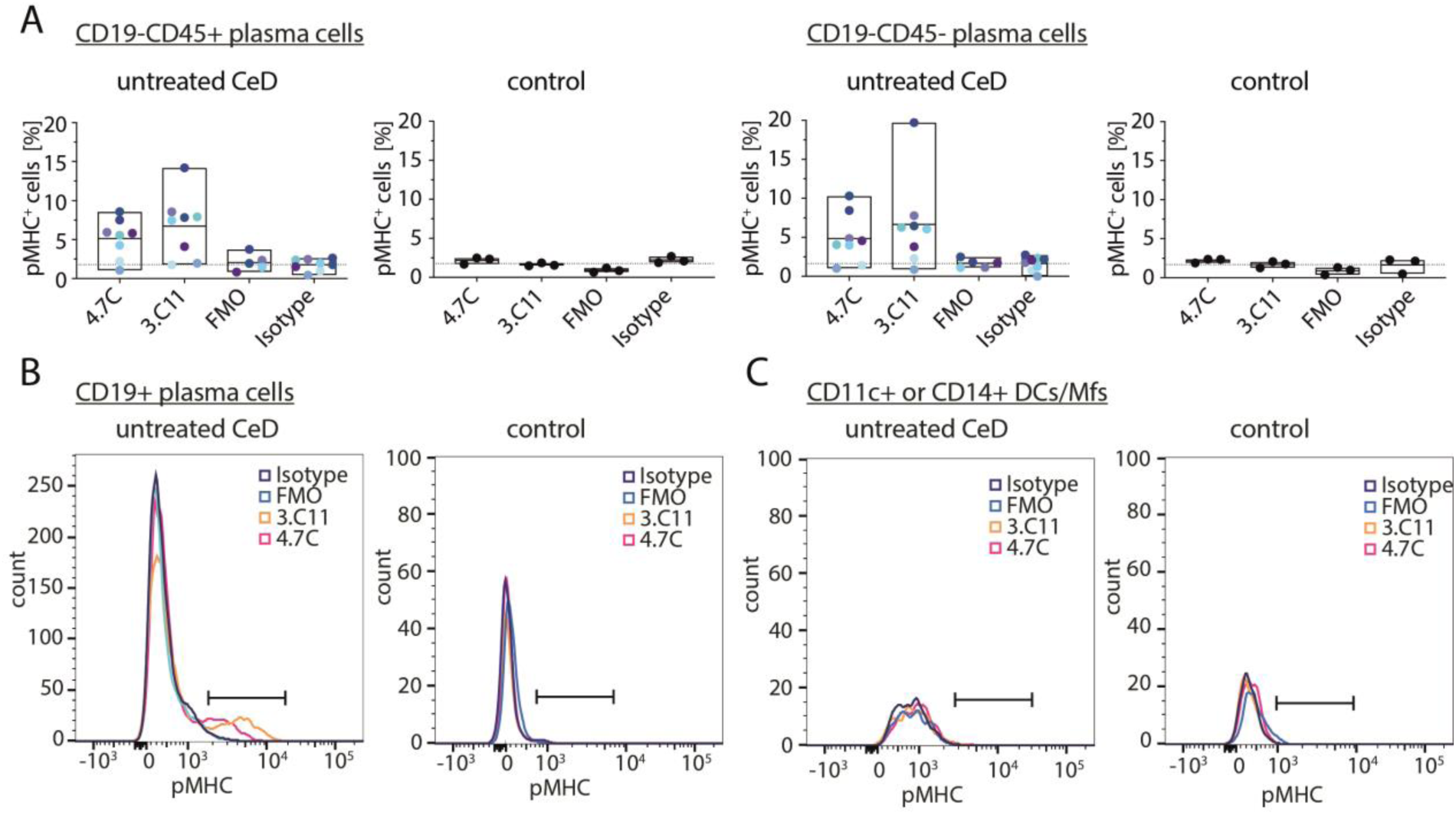
The pMHC-specific antibodies detect gluten peptide presentation on PC subsets in CeD small intestinal biopsies. Single-cell suspensions were prepared from either untreated HLA-DQ2.5^+^ CeD patients or controls with a healthy mucosal histology. Bound pMHC-specific mIgG2b antibodies were detected with an Alexa-546-conjugated secondary antibody. **A+B:** PCs were gated as live, large lymphocytes, CD3^−^CD11c^−^CD14^−^CD38^+^CD27^+^ PCs and separated into different subsets based on CD19 and CD45 expression. The frequency of pMHC^+^ cells was assessed using the TCR-like antibodies and compared to use of an isotype control antibody (isotype). Secondary antibody only (FMO) was included as a control. A. Fraction of pMHC^+^ CD45+ and CD45-PCs in untreated CeD patients (n=8) and controls (n=3). Dotted lines represent background staining using the isotype control on CeD biopsy material. **B:** Representative histograms showing staining of CD19+ PCs from an untreated CeD patient and a control subject with normal mucosal immunology. **C:** DCs/Mfs were gated as CD3^−^CD19^−^CD27^−^CD38^−^CD11c^+^CD14^+^ cells. Notably, the number of DCs/Mfs was low in all samples. Horizontal lines indicate gates set to determine fraction of pMHC^+^ cells.

**S. 8:**
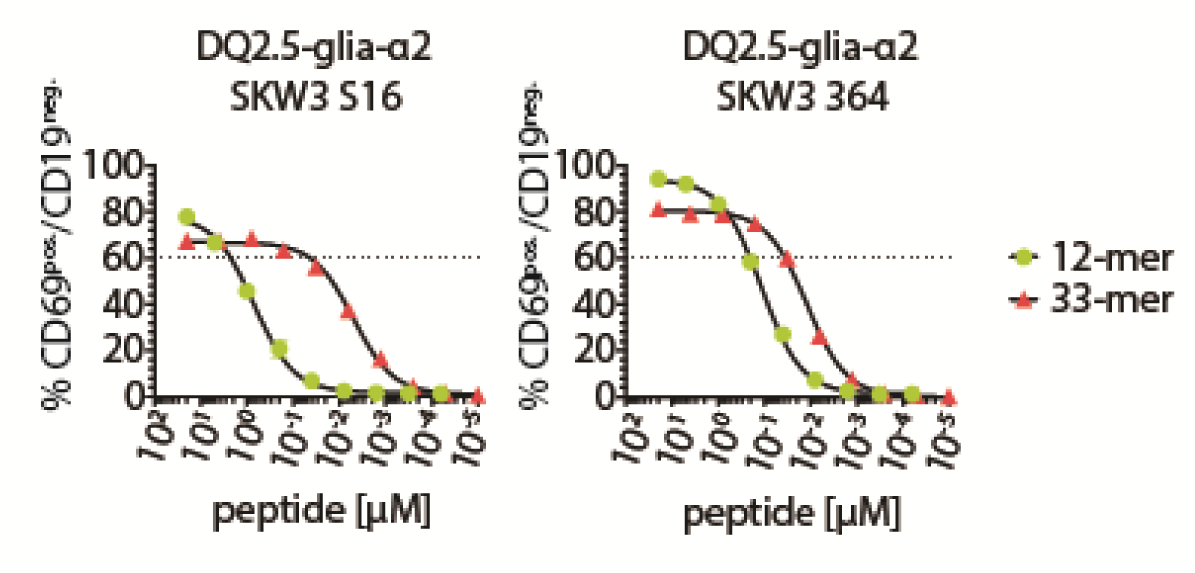
Activation of gliadin-specific SKW3 T cells. Raji B cells were loaded with a serial dilution of peptide and co-cultured with engineered gliadin-specific SKW3 T cells. T-cell activation was measured as CD69^+^ CD19^−^ cells in flow cytometry. Error bars illustrate mean ± SD of duplicates (n=2).

**Supplementary Table 1:** Design strategy for targeted libraries.

| **library name** | **randomization** | **primary transformants** | **Selection branch** |
| --- | --- | --- | --- |
| **107 H1<sup>1</sup>** | CDR H1 randomization<br>length increased by 1 or 2 residues | 3.6E9 | competition |
| **107 H3<sup>1</sup>** | CDR H3 randomization<br>fully random or retained W100 | 3.2E9 | competition |
| **107 CDR<sup>2</sup>** | 107 H1 + 107 H3 | 6.8E9 | thermal |
| **206 H1<sup>1</sup>** | CDR H1 randomization<br>length increased by 2 or 3 residues | 4.1E9 | competition |
| **206 H3<sup>1</sup>** | CDR H3 randomization<br>length increased by 1 or 2 residues | 1.2E9 | competition |
| **206 CDR<sup>2</sup>** | 206 H1 + 206 H3 | 5.3E9 | thermal |

<sup>1</sup> Libraries were designed to introduce new or improved interactions between the CDR1 and CDR3 loops of the heavy chain and the target pMHC using degenerate NNK oligonucleotides.
<sup>2</sup> Additional “107 CDR” and “206 CDR” libraries were made that contained a blend of CDR H1 and CDR H3 loop mutants.

**Supplementary Table 2.** Kinetics and affinity of affinity matured variants.

|  | Kinetics and affinity |  |  |  | Steady state<sup>c</sup> |  |
| --- | --- | --- | --- | --- | --- | --- |
| Clone | $k_{on}$ (M<sup>-1</sup>s<sup>-1</sup>) | $k_{off}$ (s<sup>-1</sup>) | $K_D$ (M) | SE $K_D$ (M) | $K_D$ (M) | SE $K_D$ (M) |
| **HLA-DQ2.5:DQ2.5-glia-α1a binders** |  |  |  |  |  |  |
| scFv 107<sup>a</sup> | 2.89x10<sup>5</sup> | 0.02151 | 7.43x10<sup>-8</sup> | 7.44x10<sup>-8</sup> | 6.70x10<sup>-8</sup> | 1.30x10<sup>-8</sup> |
| Fab 107<sup>a</sup> | 2.24x10<sup>5</sup> | 0.01601 | 7.14x10<sup>-8</sup> | 5.54x10<sup>-8</sup> | 7.41x10<sup>-8</sup> | 1.20x10<sup>-9</sup> |
| Fab 107<sup>b</sup> | 2.377x10<sup>5</sup> | 0.01698 | 7.145x10<sup>-8</sup> |  | NA | NA |
| Fab 4.7C<sup>b</sup> | 3.776x10<sup>5</sup> | 9.819x10<sup>-5</sup> | 2.600x10<sup>-10</sup> |  | NA | NA |
| Fab 4.5D<sup>b</sup> | 3.830x10<sup>5</sup> | 1.657x10<sup>-3</sup> | 4.327x10<sup>-9</sup> |  | NA | NA |
| Fab 4.6C<sup>b</sup> | 6.972x10<sup>5</sup> | 2.617x10<sup>-4</sup> | 3.754x10<sup>-10</sup> |  | NA | NA |
| Fab 4.6D<sup>b</sup> | 4.638x10<sup>5</sup> | 5.413x10<sup>-4</sup> | 1.167x10<sup>-9</sup> |  | NA | NA |
| Fab 5.6A<sup>b</sup> | 1.002x10<sup>6</sup> | 3.442x10<sup>-3</sup> | 3.436x10<sup>-9</sup> |  | NA | NA |
| Fab 15.6A<sup>b</sup> | 5.883x10<sup>5</sup> | 2.209x10<sup>-3</sup> | 3.755x10<sup>-9</sup> |  | NA | NA |
| Fab 4.7C | 9.007x10<sup>5</sup> | 1.770x10<sup>-4</sup> | 1.966x10<sup>-10</sup> | 5.40x10<sup>-7</sup> | NA | NA |
| Fab 4.7C | 6.697x10<sup>5</sup> | 9.685x10<sup>-5</sup> | 1.446x10<sup>-10</sup> | 5.60x10<sup>-7</sup> | NA | NA |
| **HLA-DQ2.5:DQ2.5-glia-α2 binders** |  |  |  |  |  |  |
| Fab 206<sup>a</sup> | 1.02x10<sup>6</sup> | 0.2291 | 2.24x10<sup>-7</sup> | 2.25x10<sup>-7</sup> | 2.54x10<sup>-7</sup> | 2.70x10<sup>-9</sup> |
| Fab 206<sup>b</sup> | 1.077x10<sup>6</sup> | 0.2462 | 2.280x10<sup>-7</sup> |  | NA | NA |
| Fab 2.B11<sup>b</sup> | 2.208x10<sup>6</sup> | 3.285x10<sup>-4</sup> | 1.488x10<sup>-10</sup> |  | NA | NA |
| Fab 3.C11<sup>b</sup> | 1.669x10<sup>6</sup> | 6.967x10<sup>-6</sup> | 4.174x10<sup>-11</sup> |  | NA | NA |
| Fab 3.C7<sup>b</sup> | 2.006x10<sup>6</sup> | 7.111x10<sup>-4</sup> | 3.544x10<sup>-10</sup> |  | NA | NA |
| Fab 3.D8<sup>b</sup> | 1.545x10<sup>6</sup> | 1.729x10<sup>-3</sup> | 1.119x10<sup>-9</sup> |  | NA | NA |
| Fab 3.F6<sup>b</sup> | 8.096x10<sup>5</sup> | 1.225x10<sup>-3</sup> | 1.513x10<sup>-9</sup> |  | NA | NA |
| Fab 12.F6<sup>b</sup> | 8.039x10<sup>5</sup> | 7.456x10<sup>-4</sup> | 9.275x10<sup>-10</sup> |  | NA | NA |
| Fab 3.C11 | 1.190x10<sup>6</sup> | 1.126x10<sup>-4</sup> | 9.462x10<sup>-11</sup> | 3.10x10<sup>-7</sup> | NA | NA |
| Fab 3.C11 | 1.387x10<sup>6</sup> | 1.118x10<sup>-4</sup> | 8.064x10<sup>-11</sup> | 1.30x10<sup>-7</sup> | NA | NA |
| Fab 3.C11 | 1.335x10<sup>6</sup> | 1.206x10<sup>-4</sup> | 9.039x10<sup>-11</sup> | 1.20x10<sup>-7</sup> | NA | NA |

Kinetics were determined by fitting data to a 1:1 Langmuir binding model.
<sup>a</sup> Determined from single cycle kinetics.
<sup>b</sup> Determined from one concentration of protein in off-rate screening. $K_D$ s obtained with this method disregarded when calculating average values.
<sup>c</sup> Steady state $K_D$ was derived from the single cycle kinetics runs.
NA, not available.

**Supplementary Table 3.**
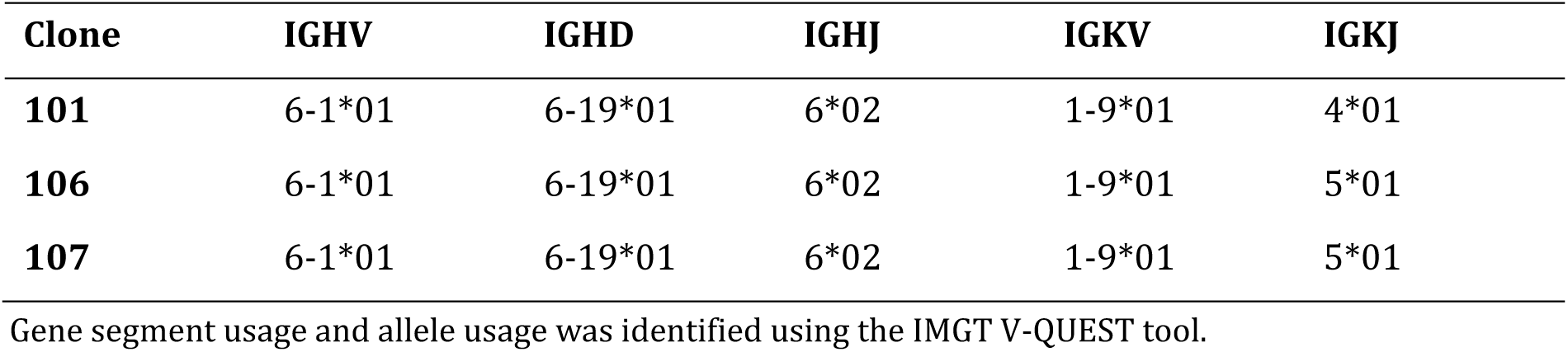
Gene segment usage among the HLA-DQ2.5:DQ2.5-glia-α1a-specific clones (first generation).

| Clone | IGHV | IGHD | IGHJ | IGKV | IGKJ |
| --- | --- | --- | --- | --- | --- |
| 101 | 6-1*01 | 6-19*01 | 6*02 | 1-9*01 | 4*01 |
| 106 | 6-1*01 | 6-19*01 | 6*02 | 1-9*01 | 5*01 |
| 107 | 6-1*01 | 6-19*01 | 6*02 | 1-9*01 | 5*01 |

Gene segment usage and allele usage was identified using the IMGT V-QUEST tool.

**Supplementary Table 4.** Antibodies used for flow cytometric analyses of peptide-presenting cells in human intestinal samples.

| **Antigen** | **Conjugate** | **Clone** | **Supplier** | **Dilution** |
| --- | --- | --- | --- | --- |
| mIgG2b 4.7C/3.C11 | - | 4.7C, 3.C11 | In-house | 10 µg/ml |
| Isotype control | - | OMV | In-house | 10 µg/ml |
| mIgG | Alexa-546 | Polyclonal | Invitrogen | 1 µg/ml |
| CD3 | FITC | OKT3 | Biolegend | 1:20 |
| CD11c | APC | S-HCl-3 | BD Biosciences | 1:20 |
| CD14 | APC | HCD14 | Biolegend | 1:20 |
| CD19 | PE-Cy7 | HIB19 | Biolegend | 1:20 |
| CD27 | BV421 | O323 | Biolegend | 1:20 |
| CD38 | APC-Cy7 | HIT2 | Biolegend | 1:20 |
| CD45 | BV510 | H130 | Biolegend | 1:20 |

